# Mesenchymal stem cells (MSCs) offer a drug tolerant and immune- privileged niche to *Mycobacterium tuberculosis*

**DOI:** 10.1101/521609

**Authors:** Neharika Jain, Haroon Kalam, Lakshyaveer Singh, Vartika Sharma, Saurabh Kedia, Prasenjit Das, Vineet Ahuja, Dhiraj Kumar

## Abstract

Anti-tuberculosis (TB) drugs while being highly potent *in vitro* require prolonged treatment to control *Mycobacterium tuberculosis* (*Mtb*) infections *in vivo*. We report here, mesenchymal stem cells (MSCs) shelter *Mtb* to help tolerate anti-TB drugs. MSCs uptake *Mtb* readily and allow them grow unabated despite having functional innate pathway of phagosome maturation. Unlike macrophage-resident ones, MSC-resident *Mtb* tolerates anti-TB drugs remarkably well, a phenomenon requiring proteins ABCC1, ABCG2 and vacuolar-type H^+^ATPases. Additionally, contrary to what is classically known, IFNγ and TNFα aid mycobacterial growth within MSCs. Mechanistically, evading drugs and inflammatory cytokines by MSC-resident *Mtb* is dependent on elevated PGE2 signaling, which we subsequently verified *in vivo* analyzing sorted CD45^-^CD73^+^SCA1^+^-MSCs from the lungs of infected mice. Moreover granulomas from human pulmonary and extra-pulmonary TB show presence of MSCs co-inhabiting with *Mtb*. Together the results show targeting the immune-privileged niche, provided by MSCs to *Mtb*, can revolutionize tuberculosis prevention and cure.

## Introduction

*Mycobacterium tuberculosis* (*Mtb*) continues to infect, cause illness (tuberculosis) and kill large number of individuals globally. Among numerous factors that thwart any tuberculosis control program, lack of an effective vaccine and long duration of treatment are the two most critical ones^1,2^. Long treatment duration is majorly attributed behind non-compliance and emergence of drug-resistant tuberculosis including multi- and extensively drug-resistant (MDR and XDRs respectively) ones^1^. While standard-of-care anti-TB drugs are very efficient in killing *Mtb* in liquid culture and during *ex vivo* infection studies in macrophages, their efficacy is dramatically compromised during *in vivo* infection studies and in the clinical practices, requiring prolonged treatment duration^3^. It is believed that *Mtb* undergoes metabolic adaptations within host granulomas, which render these bacteria less vulnerable to the standard drugs ^4,5^. Driving factors, which cause such adaptations include nitric oxide (NO), redox stress (ROS), low oxygen (hypoxia), low nutrients or altered carbon source ^4,6-11^.

Curiously, whatever we know about the intracellular lifestyle of mycobacteria in the hosts is mostly through studies on macrophages ^12,13^. Are there additional niches of mycobacteria *in vivo*, which could facilitate the perceived metabolic adaptations? While there is no clear answer to the above assumption, there are certainly different other cell types which get infected inside the host including lung epithelial cells, macrophages, neutrophils, dendritic cells, adipocytes and mesenchymal stem cells (MSCs) ^14-18^. MSCs are peculiar among these cells since they were first reported to dampen the host immunity against tuberculosis around the granulomas ^19^. Subsequently it was observed that these cells are the site of persistent or latent bacterial infection ^20^. Interestingly, latent bacteria are perceived to be more tolerant to anti-TB drugs ^21-23^. Moreover, MSCs are classically known for their immune-modulatory functions ^24-26^. Whether MSCs provide a privileged niche to mycobacteria allowing them to withstand drug and evade host immunity remains unclear. Potential benefits mycobacteria enjoys within these cells continue to remain obscure due to lack of systematic studies on the intracellular lifestyle of *Mtb* within MSCs. In this study, using adipose tissue-derived mesenchymal stem cells (ADSCs), we show that *Mtb* not only escapes the effect of anti-TB drugs while residing within ADSCs but also effectively evade host immune mediators. We further establish the mechanism behind these unusual properties of ADSCs and show their relevance during *in vivo* infection in mice as well as studies on the human subjects.

## Results

### Adipose-derived Mesenchymal stem cells (ADSCs) support mycobacterial growth

Human primary adipose-derived mesenchymal stem cells obtained commercially were first characterized for expression of cell-surface markers like CD73 and CD271 as well as their ability to differentiate into three different lineages i.e. adipocytes, chondrocytes and osteocytes (Fig. S1A and S1B). We infected ADSCs with *GFP-H37Rv* (MOI 1:10) with ∼80 percent efficiency (Fig. 1A). Mean fluorescence intensity (MFI) measurements at 6 days post-infection showed that *Mtb* within ADSCs multiplied well (Fig. 1B), which we also confirmed by colony forming unit (CFU) counts upon plating the bacteria released by lysing the infected ADSCs (Fig. 1C). A time-course growth analysis using CFU counts showed massive increase in *Mtb* CFU at 9 and 12 days post-infection (Fig. 1C), which however was not the case with the vaccine strain BCG that showed marked decline in survival within ADSCs by 3 days post-infection (Fig. 1D). While *H37Rv* survived well in human primary macrophages (Fig. 1E) and within THP-1 derived macrophages (Fig. 1F), consistent with previous reports from several groups including ours ^27-29^, its multiplication within macrophages was markedly subdued when compared with that observed within ADSCs (Fig. 1). Infection with *H37Rv* did not result in spontaneous differentiation of ADSCs to any of the three lineages mentioned above (Fig. S1C). A microarray analysis of ADSCs infected with *H37Rv* for 6 days also did not reveal any significant change in expression of genes involved in differentiation into adipocyte, chondrocyte or osteocyte (Fig. S1D-E and Table S1). Microarray analysis showed significant regulation of genes belonging to usual functional classes like immune regulation, inflammation, response to stress, transport pathways and cholesterol metabolism etc. (Fig S1F).

**Figure 1:**
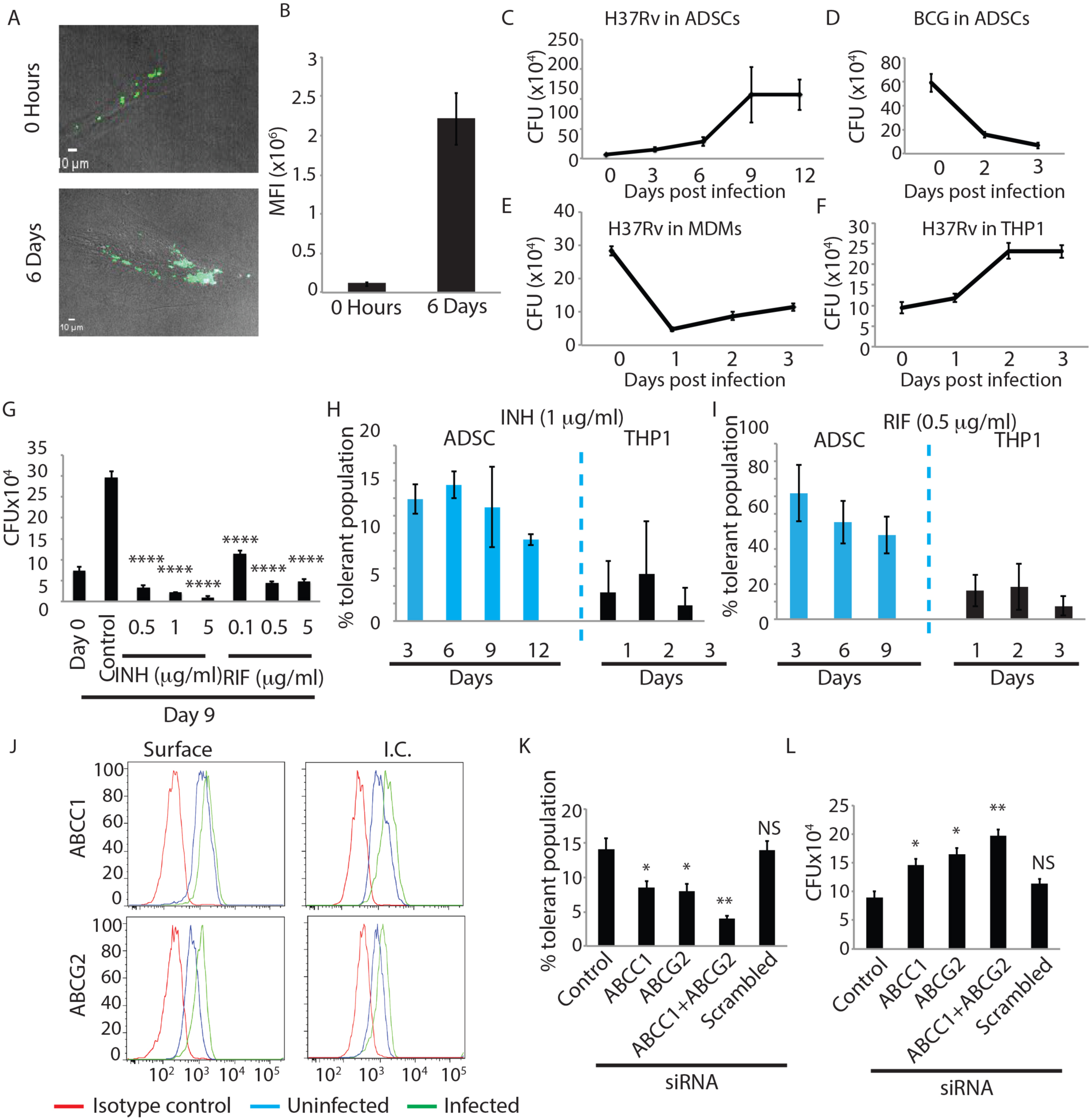
ADSCs support better H37Rv survival and high drug tolerance. **(A)** ADSCs were infected with GFP-Rv at 1:10 MOI, representative confocal images at 0 hour and 6^th^ day post infection are shown. **(B)** MFI of GFP-Rv per cell calculated across 10 fields repeated in more than triplicate. **(C)** Growth kinetics of H37Rv within ADSCs across 12 days post-infection. **(D)** Growth kinetics of vaccine strain BCG within ADSCs on 2 and 3 days post-infection. **(E)** Growth kinetics of H37Rv within human MDMs and **(F)** in THP-1 macrophages across 0, 1,2 and 3 days post-infection. **(G)** H37Rv infected ADSCs were treated with isoniazid (INH), and rifampicin (RIF), 0.1 – 5 µg/ml, for 24 hours on 8^th^ day post-infection, and cells were plated for CFU enumeration. **(H)** Percent drug tolerant bacterial population to INH (1µg/ml) within infected ADSCs (blue) on 3^rd^, 6^th^, 9^th^, 12^th^ day and within THP-1 macrophages (black) on 1^st^, 2^nd^ and 3^rd^ days post-infection, respectively. **(I)** Percent drug tolerant bacterial population to RIF (0.5 µg/ml) within infected ADSCs (blue) on 3^rd^, 6^th^, 9^th^ days and within THP-1 macrophages (black) on 1^st^, 2^nd^ and 3^rd^ days post-infection, respectively. **(J)** Line histogram of surface and intracellular (I.C.) staining of ABCC1/MRP-1 and ABCG2/BCRP in uninfected (Blue line) and in H37Rv-infected ADSCs (Green line), 6 days post-infection. Red line represents the isotype control. **(K)** Percent INH (1µg/ml) tolerant bacterial population in ADSCs after knocking down ABCC1 (200 nM siRNA) or ABCG2 (200 nM siRNA) alone or in combination, in H37Rv-infected ADSCs. The siRNAs were added 24 hours prior to treatment with INH for additional 24 hours and CFU plating was carried out on 6^th^ day post-infection. **(L)** H37Rv CFU from ADSCs on 6^th^ day after siRNA mediated knockdown of ABCC1 and ABCG2 for 48 hours prior to the time point. Data are from three or more independent experiments. Error bar represent S.E.M. *p < 0.05, **p < 0.005, ***p < 0.0005, ****p < 0.00005, NS ‘not significant’ by two-tailed Student’s t-test. Scale bars, 10 µm.

### ADSC resident *Mycobacterium tuberculosis* shows drug tolerant phenotype

Since MSCs were reported to serve as a site for bacterial persistence ^20^ we were keen to understand how *Mtb* residing within these cells responds to anti-TB drugs. We treated *Mtb*-infected ADSCs with different doses of isoniazid (INH) or rifampicin (RIF) for 24 hours before harvesting the cells and CFU plating on 9th day post-infection. Even at doses as high as 5µg/ml for INH, ∼10% of *Mtb* tolerated the drug (Fig. 1G). The percent tolerant population was more than 15% at 0.5 µg/ml as well as at 1 µg/ml of INH (Fig. 1G). In case of RIF, nearly 50% of bacteria were tolerant to the drug at 0.1 µg/ml, which did not go down below 15% even at doses as high as 5µg/ml (Fig. 1G). Interestingly, drug tolerant phenotype of ADSC-resident *Mtb* was independent of time spent within ADSCs as drug (INH or RIF) tolerant *H37Rv* were observed as early as 3 days post-infection and maintained at 6, 9 and 12 days post-infection (Fig. 1H and 1I). At similar doses and for similar duration of treatment (i.e. 24 hours) within macrophages, there were hardly any surviving bacteria in case of INH (∼2-4%) while there were nearly 10-15% tolerant bacteria in case of RIF (Fig. 1H and 1I). Thus it was evident that ADSCs provide an environment, which allowed *Mtb* to tolerate anti-TB drugs.

### Host ABC transporters ABCC1 and ABCG2 play key role in bacterial drug tolerance

MSCs are known to express high level of ABC family transporters or efflux pumps, which are often attributed to drug tolerance in case of cancer ^30,31^. In our microarray data we did observe slight but consistent change in the expression of several of the ABC transporters including ABCC1 and ABCG2, which are also known as MRP1 and BCRP respectively ^32,33^ (Table S1). In fact, ADSCs showed increase in expression of ABCC1 and ABCG2 upon *Mtb* infection in MOI dependent manner (Fig S2A). Both intracellular as well as surface expression of ABCC1 and ABCG2 were higher in H37Rv infected ADSCs with respect to the control cells (Fig. 1J). To test whether ABCC1 and ABCG2 were involved in imparting drug tolerance, we used known pharmacological inhibitors against them. Treatment with novobiocin (an ABCG2 inhibitor) or with MK571 (ABCC1 inhibitor) led to a decline in the drug tolerant *Mtb* population (Fig. S2B-C). Novobiocin however is also a well-known DNA gyrase inhibitor and it could actually kill *Mtb* even *in vitro* in liquid cultures (Fig S2D). Unlike novobiocin, MK571 treatment did not have any effect on *Mtb* growth *in vitro* (Fig. S2E). Since pharmacological inhibitors may still have off-target effects, we knocked down these transporters using specific siRNAs. Knocking down either ABCC1 or ABCG2 led to a substantial decline in the drug-tolerant bacterial population within ADSCs (Fig. 1K and Fig. S2F). When ABCC1 and ABCG2 both were knocked down simultaneously, the tolerant bacterial population was almost wiped out reaching nearly 2-4 % (Fig. 1K). There was no such effect on drug-tolerant population when scrambled siRNA were used (Fig. 1K). While the results with tolerant population did indicate the role of ABCC1 and ABCG2 in *Mtb* drug tolerance within ADSCs, in parallel experimental groups where no drug was used, knocking down ABCC1 and ABCG2 led to a considerable increase in bacterial CFU (Fig. 1L). The increase in bacterial CFU was higher when both ABCC1 and ABCG2 were simultaneously knocked down whereas there was no effect when scrambled siRNA was used as control (Fig. 1L). Similar results were also obtained when ABCC1 or ABCG2 were inhibited by corresponding pharmacological inhibitors (Fig. S2G-H).

### Role of lysosomal function in mycobacterial drug tolerance in ADSCs

While ABCC1 and ABCG2 seemed important for drug tolerance, their role in bacterial killing as evident in figure 1L was surprising. Curiously, the role of ABCC1 and ABCG2 seemed more to do with the lysosomal function rather than actual efflux activity at the cell surface since inhibition of vacuolar type H^+^ ATPases by bafilomycin A1 (BafA1) also completely wiped out drug tolerant *Mtb* within ADSCs (Fig. 2A). Another lysosomal acidification inhibitor choloroquine (CQ) had similar effect (Fig. 2A). However conditions, which led to increased lysosomal maturation like rapamycin treatment showed drug tolerant population at par with the control ADSCs (Fig 2A). Since rapamycin is a well-known inducer of autophagy ^34^, we also verified it using autophagy inhibitor 3-methyladenine (3MA), which expectedly led to a decline in the drug tolerant population (Fig. 2A). Thus lysosomal function was probably important to achieve drug tolerant phenotype within ADSCs. Interestingly, in the absence of INH, conditions, which resulted in reducing the drug tolerant population, helped bacterial survival. Thus, BafA1, 3MA and CQ treatment resulted in increased bacterial CFU whereas rapamycin treatment led to a decline in the CFU suggesting role of lysosomal killing mechanism in MSCs (Fig. 2B). Before further exploring into the mechanism of drug tolerance within ADSCs, we wanted to compare this phenomenon with the reported instances of drug tolerance in macrophages ^35^.

**Figure 2:**
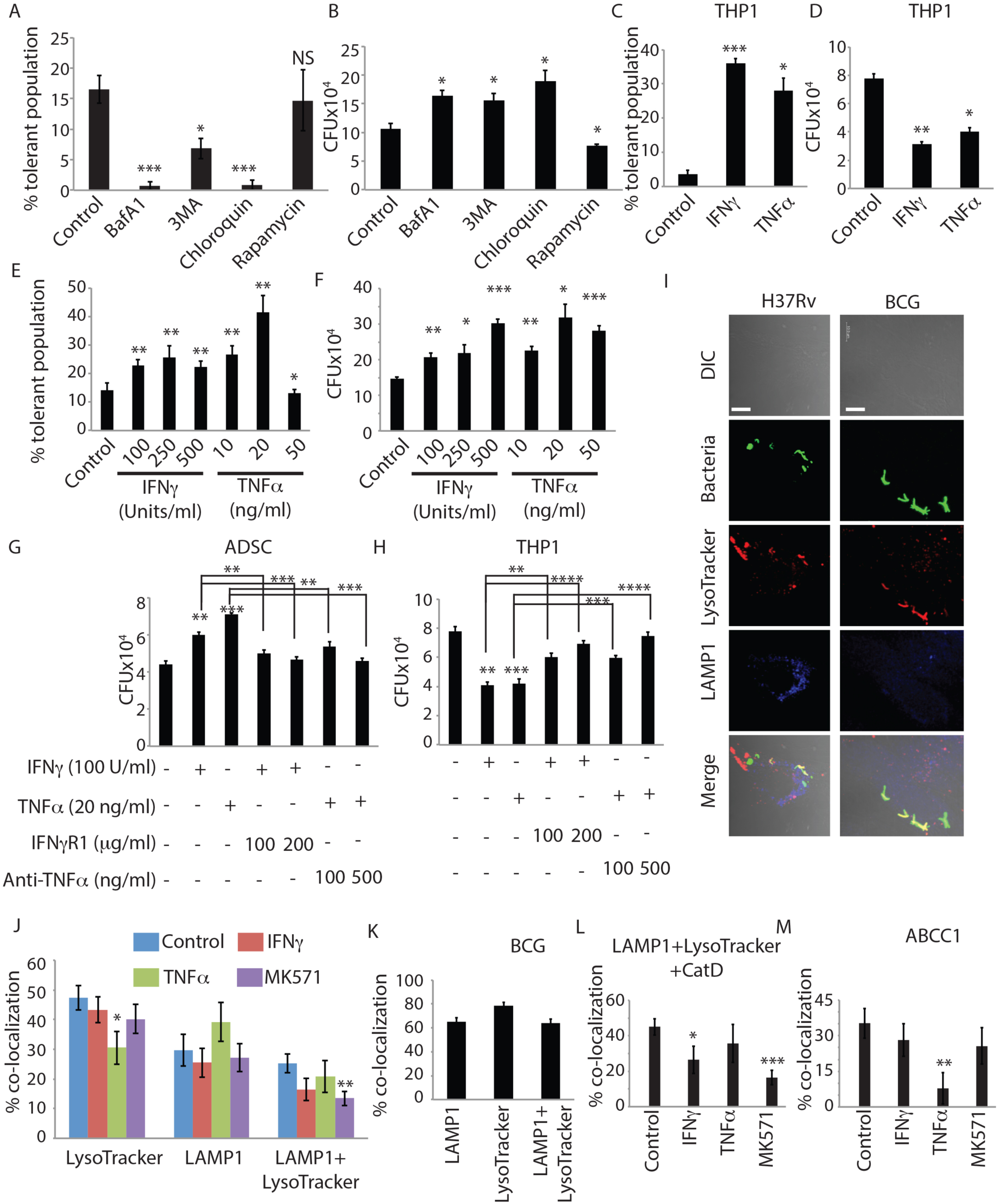
Lysosomal killing of bacteria in MSCs and effect of inflammatory cytokines. **(A)** Percent INH tolerant H37Rv population after addition of autophagy modulators bafilomycinA1 (BafA1, 100 nM), 3-methyl adenine (3-MA, 5mM), chloroquine (100 µM), and rapamycin (100 nM) for 6 hours before CFU plating on 6^th^ day post-infection in ADSCs. **(B)** H37Rv infected ADSCs were treated with BafA1 (100nM), 3-MA (5mM), chloroquine (100 µM) and rapamycin (100nM) for 6 hours prior to CFU plating on the 6^th^ day post-infection. **(C)** Percent INH-tolerant H37Rv population after treatment of infected THP-1 macrophages with 100 units/ml IFNγ and 20 ng/ml TNFα for 24 hour before CFU plating on 3^rd^ day post-infection. **(D)** THP-1 macrophages were infected with H37Rv and treated with 100 units/ml IFNγ or 20 ng/ml TNFα for 24 hours prior to CFU plating on the 3^rd^ day post-infection. **(E)** Dose dependent effect of IFNγ (100, 250,500 units/ml) and TNFα (10, 20, 50 ng/ml) treatment (24 hours each) on percent INH tolerant population in H37Rv-infected ADSCs. **(F)** H37Rv survival within ADSCs after treatment with increasing doses of IFNγ or TNFα for 24 hours prior to CFU plating on 6^th^ day post-infection. **(G, H)** CFU assay of infected ADSCs (G) and infected THP-1 macrophages (H) after 24 hours of IFNγ or TNFα treatment in the presence of different doses of IFNγR1 and anti-TNFα purified proteins respectively. IFNγR1 or anti-TNFα was added along with IFNγ or TNFα treatments respectively **(I)** Representative confocal images of PKH67-labeled H37Rv and BCG infected ADSCs on 3^rd^ day post-infection, co-stained with LysoTracker Red and LAMP-1 antibody. **(J)** Percent colocalization of PKH67 labeled H37Rv with compartments stained for LysoTracker Red, LAMP-1 or LysoTracker+LAMP1 in ADSCs which were either untreated control or treated with IFNγ (100 units/ml), TNFα (20 ng/ml) or MK571 (50 µM) for 24 hours prior to fixing the samples at 3^rd^ day post-infection. **(J)** Percent colocalization of BCG with LysoTracker Red, LAMP-1 and LAMP-1+LysoTracker compartments at 3 days post-infection in ADSCs. **(L)** Colocalization of H37Rv to Cat D, LAMP-1 and lysoTracker Red triple positive compartment in infected ADSCs which were either untreated or treated with IFNγ (100 units/ml), TNFα (20 ng/ml) or MK571 (50 µM) for 24 hours prior to processing the samples on 3^rd^ day post-infection. **(M)** Percent colocalization of H37Rv with ABCC1 within ADSCs in untreated or upon treatment with IFNγ (100 units/ml), TNFα (20 ng/ml) or MK571 (50 µM) for 24 hours prior to harvesting the samples on 3^rd^ day post-infection. Error bar represent S.E.M. *P < 0.05, **P < 0.005, ***P < 0.0005, ****P < 0.00005, NS ‘not significant’ by two-tailed Student’s t-test. Scale bars, 10 µm. All the data is compiled from three independent experiments.

### Effect of inflammatory cytokines IFNγ and TNFα on drug tolerance within ADSCs

In macrophages, activation with inflammatory cytokines is known to induce drug tolerant phenotype of *Mtb* ^35^. We reconfirmed, in THP-1 macrophages, treatment with IFNγ or TNFα led to a substantial increase in drug tolerant population from ∼3-4% in control to 30-40% in the activated cells (Fig. 2C). At similar doses of these cytokines, in the absence of drug, nearly 50% of the bacteria got killed, in agreement with the anti-bacterial state these cytokines impart to the activated macrophages ^35^. In ADSCs, IFNγ treatment at 100, 250 and 500 units/ml led to almost dose dependent increase in the INH tolerant population (Fig. 2E). In case of TNFα treatment, INH tolerant *Mtb* population was higher at 10 and 20 ng/ml, which came down to almost control levels at 50ng/ml (Fig. 2E). TNFα at 20 ng/ml was much more potent in inducing drug tolerance than IFNγ at any doses studied. More startling observation however was the case where *Mtb*-infected ADSCs were treated with these cytokines in the absence of drugs. There was nearly dose dependent increase in bacterial CFU upon treatment of *Mtb*-infected ADSCs with IFNγ or TNFα (Fig. 2F). The pro-bacterial effect of IFNγ and TNFα on *Mtb*-infected ADSCs was specific to the stimulus and corresponding downstream signaling since upon neutralization with purified IFNγR1 or with anti-TNFα antibody, we could revert the pro-bacterial effect of IFNγ or TNFα stimulation on bacterial CFU (Fig. 2G). Expectedly, with the similar neutralization study in THP-1-derived macrophages, there was rescue of *Mtb* from cytokine-mediated killing (Fig. 2H). Thus *Mtb*-infected macrophages and MSCs respond in contrasting manner to IFNγ and TNFα stimulus.

### Analysis of intracellular niches of *Mtb* shows classic phagosome maturation dynamics in ADSCs

Results from conditions like ABCC1 or ABCG2 knockdown, BafA1 treatment or IFNγ or TNFα treatments, all of which led to increase in bacterial survival suggest that ADSCs, despite supporting robust growth of the bacteria, keep actively killing the bacilli. To check whether phagosome maturation pathways as observed in macrophages are operational in similar fashion during *Mtb* infection in ADSCs, we assayed for *Mtb* co-localization with early phagosomes (RAB5), late phagosomes (RAB7), lysosomes (LAMP1) and acidified lysosomes (LAMP1 and LysoTracker). At any given time post-infection, a large number of bacteria (∼40-50%) stayed within RAB5 positive early phagosomes inside the ADSCs (Fig. S3A). While only 2-5% bacteria were ever present in RAB7 positive late phagosomes (Fig. S3B). Similar distribution of *Mtb* is also reported within macrophages as reported by others and also confirmed by us ^36^ (Fig. S3C). This reflects the phagosome maturation arrest inflicted by *Mtb* in the infected macrophages ^36,37^. However unlike macrophages where LAMP1-*Mtb* or Lysotracker-*Mtb* co-localization rarely crosses ∼15% ^27^, there are more bacteria (∼30-40%) present in LAMP1 or Lysotracker-LAMP1 double positive compartments in ADSCs (Fig. 2I-J, S3D). The matured lysosomes, i.e. LAMP1 positive acidified compartments indeed reflect the killing mechanism in ADSCs since ∼80% of intracellular BCG, the strain that gets killed within ADSCs, are present in Lysotracker-LAMP1 double positive compartments (Fig. 2K). Interestingly, treatment with IFNγ, TNFα or MK571 in general led to a decline in bacterial localization to LysoTracker or LAMP1+LysoTracker compartments however LAMP1 compartment alone showed only marginal decline (Fig. 2J, S3D). Exclusion of Cathepsin D, the lysosomal protease, from LAMP1-Lysotracker positive compartment strongly correlated with increased bacterial survival upon IFNγ or MK571 treated cells (Fig. 2L, S3D). Interestingly, ABCC1 and ABCG2 were also found to co-localize with *Mtb* suggesting their recruitment to phagosomes (Fig. S3E). At least in case of TNFα treatment, exclusion of ABCC1 from the LAMP1-LysoTracker compartment was highly significant whereas IFNγ or MK571 treatment showed only marginal decline (Fig. 2M) leaving an impression that ABCC1 is probably directly involved in bacterial killing. However their direct role in *Mtb* killing is supported by only one set of evidences-increased bacterial survival upon their knockdown or inhibition. The strong correlation between bacterial killing and their co-localization to LAMP1-Lysotracker-CatD compartment nonetheless suggest active role of classic phagosome maturation pathways in bacterial killing with ADSCs.

Interestingly, in contrast to what is known in macrophages^27^, *Mtb* very rarely co-localized to autophagosomes and xenophagy flux was completely absent in ADSCs (Fig S4A), suggesting little role if any, of autophagy in controlling *Mtb* within ADSCs. Curiously, ADSCs showed very high basal autophagy flux (Fig. S4B), which presumably is critical for maintaining the stem cell like property of these cells, highlighting the segregation of homeostatic and anti-bacterial arms of autophagy as reported by us earlier ^38^. Moreover treatment with IFNγ led to increased autophagy flux in ADSCs, unlike what was reported previously for macrophages (Fig. S4C) ^39^. Unlike macrophages, IFNγ treatment had no effect on cellular ROS generation in ADSCs (Fig. S4D). To some extent, this could explain why IFNγ failed to induce killing of *Mtb* within ADSCs.

### The lipid mediator PGE2 helps MSCs exhibit pro-bacterial attributes

Results so far establish that mesenchymal stem cells are uncharacteristically pro-bacterial in nature, at least during mycobacterial infections, helping them evade anti-TB drugs as well as classic host immune mediators like IFNγ and TNFα. However, there still was no clue on how MSCs execute these behaviors. To understand the mechanistic basis of the observed results, we went back to our microarray data to identify genes showing significant regulation upon *Mtb* infection in ADSCs. The anti-inflammatory as well as immune-modulatory functions of MSCs are well known, however in all such known cases, MSCs execute its role by modulating functions of other cells, including T cells and macrophages ^40-42^. Some of the key mediators that help MSCs execute these functions are PGE2, IDO1, IL6, CCL2, VEGFC, LIF etc. ^24,42-46^. In our microarray data, genes from the PGE pathway like PTGS2, PTGES and PTGR2 showed nearly 8, 4 and 4 folds (log_2_) increase in expression respectively (Fig. S5A). Similarly IDO1 showed 6 folds increase, whereas LIF, IL6, CCL2 and VEGF each showed more than 3 fold increase in expression post-infection (Fig. S5A). We first tested PGE2 levels in the culture supernatants of ADSCs that were infected with *H37Rv*. Consistent with the microarray data PGE2 ELISA confirmed increased synthesis and secretion of PGE2 from *Mtb*-infected ADSCs (Fig. 3A). Interestingly, treatment with IFNγ or TNFα further increased PGE2 levels in the culture supernatants whereas MK571 treated cells showed almost similar level of PGE2 as infected control cells (Fig. 3A). We used celecoxib, a widely used PTGS2 (or COX2) inhibitor, which is also an FDA approved drug in the market, as a negative control. *Mtb*-infected ADSCs treated with celecoxib showed negligible PGE2 levels by ELISA (Fig. 3A). Next we treated *Mtb*-infected ADSCs with celecoxib at 50, 150 and 250µM concentrations under all the conditions tested so far in this study. Treatment with celecoxib reduced *Mtb* CFU in a dose dependent manner across the conditions including infection alone or when treated with IFNγ, TNFα or MK571 (Fig. 3B). Similar results were also obtained with EP2 receptor (receptor for PGE2) antagonist PF04418948, suggesting involvement of signaling through PGE2 pathway in bacterial survival (Fig. S5B). Celecoxib was also effective in killing *Mtb* within macrophages however not as dramatically as observed in MSCs (Fig. S5C). Unlike ADSCs, there was no increase in PGE2 release by THP-1 macrophages upon infection or treatment with IFNγ or TNFα (Fig. S5D). We also verified these results by knocking down PTGS2 (COX2) using specific siRNAs (Fig. 3C and S5E). Confocal microscopy revealed that majority of bacteria in celecoxib treated cells or PF04418948 treated cells or COX2 siRNA treated cells co-localized with LAMP-1, Lysotracker as well as CatD (Fig. 3D and S5F). Interestingly, COX2 inhibition by celecoxib also helped limit the drug tolerant phenotype in ADSCs against INH, irrespective of treatment with IFNγ or TNFα (Fig. 3E). Moreover, MK571, which itself decreases drug-tolerant *Mtb* in ADSCs when combined with celecoxib treatment further decreases the drug-tolerant population of *Mtb* within ADSCs (Fig 3E). The effect of celecoxib on bacterial drug tolerance was PGE2-mediated and not due to possible role of certain COX2 inhibitors directly on bacterial drug-resistance protein MDR1 ^47^ since knocking down COX-2 also led to a remarkable decline in INH-tolerant as well as rifampicin-tolerant *Mtb* population within ADSCs (Fig. 3F and 3G respectively). Efficacy of COX2 knockdown by siRNA on bacterial drug tolerance also rule out role of PGE2 inhibitors in directly regulating the efflux proteins as reported previously ^48^.

**Figure 3:**
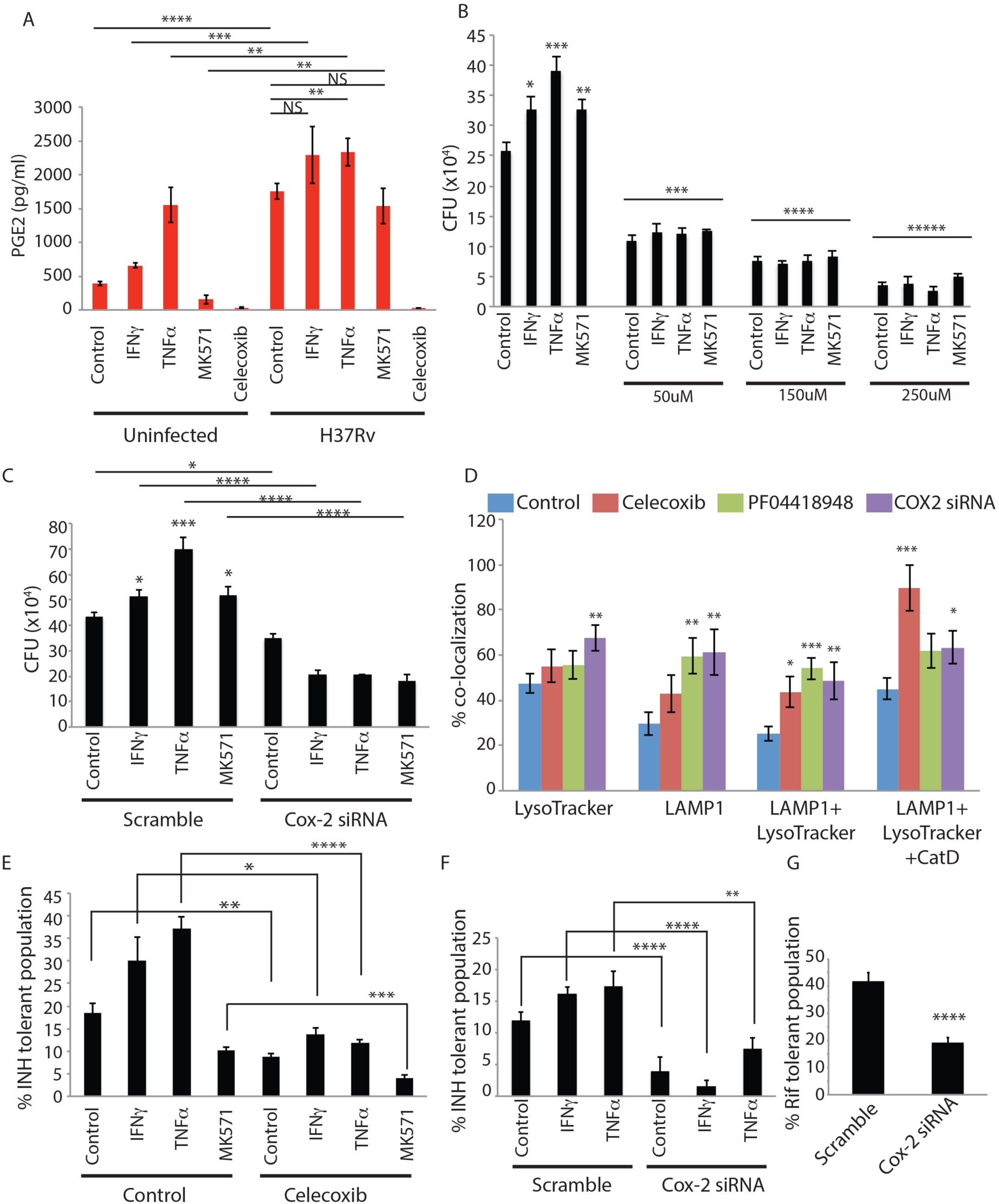
Lipid mediator PGE2 modulates lysosomal activity in *Mtb*-infected ADSCs. **(A)** Uninfected or H37Rv infected ADSCs which were either untreated or treated with IFNγ (100 units/ml), TNFα (20 ng/ml), MK571 (50 µM) and celecoxib (250 µM) for 24 hours before performing supernatant ELISA on 6^th^ day post-infection (post-seeding in uninfected cells). **(B)** H37Rv infected ADSCs were treated with different doses of celecoxib (50um, 150uM, 250uM) in addition to IFNγ (100 units/ml), TNFα (20 ng/ml) or MK571 (50 µM) for 24 hours before the cells were harvested for CFU plating on 6^th^ day post-infection. **(C)** H37Rv infected ADSCs were treated with Cox-2 siRNA or scramble siRNA (100nM siRNA) along with treatment with IFNγ (100 units/ml), TNFα (20 ng/ml) or MK571 (50 µM) for 24 hours prior to CFU plating on 6^th^ day post-infection. **(D)** Colocalization of PKH67 labeled H37Rv with LysoTracker Red, LAMP-1, and Cathepsin D stained compartments in ADSCs after 24 hours of treatment with Celecoxib (250 µM), PF04418948 (500 nM) or Cox-2 siRNA (100 nM) before harvesting samples on 3^rd^ day post-infection. **(E)** Percent INH tolerant population in ADSCs treated with celecoxib (250 µM) along with addition of IFNγ (100 units/ml), TNFα (20 ng/ml) or MK571 (50 µM) for 24 hours prior to CFU plating 6^th^ day post-infection. **(F)** Percent INH-tolerant H37Rv population after treatment of infected ADSCs with 100 units/ml IFNγ or 20 ng/ml TNFα for 24 hours in scramble or Cox-2 siRNA (100nM) treated cells and CFU plating was performed on 3^rd^ day post-infection. **(G)** Percent RIF tolerant bacterial population in infected ADSCs treated with scrambled or cox-2 siRNA (100 nM) for 24 hours prior to CFU plating on 6^th^ day post-infection. Error bar represent S.E.M. *P < 0.05, **P < 0.005, ***P < 0.0005, ****P < 0.00005, NS ‘not significant’ by two-tailed Student’s t-test.

### MSCs serve as a niche for *Mtb* during *in vivo* infection allowing drug tolerance in PGE2 dependent manner

While all the results so far were performed on human primary adipose tissue-derived mesenchymal stem cells, we next wanted to explore whether these cells actually get involved during *in vivo* infection in mice and humans as well as to know whether PGE2 signaling plays similar role *in vivo*. We infected C57BL/6 mice with *H37Rv* through aerosol challenge and 4 weeks post-infection, these animals were divided into four groups: control, celecoxib (50mg/kg), INH (10mg/kg) or INH+celecoxib; and treated for subsequent 4 and 8 weeks. From the initial bacterial load of 100-150 per animal, it reached around 10^6^ per animal by end of 4 weeks, 2×10^6^ by the end of eight weeks and 3×10^6^ by the end of 12 weeks in the lungs (Fig 4A). While celecoxib treatment alone significantly reduced bacterial CFU after 4 weeks of treatment, the effects were not visible at 8 weeks post-treatment (Fig. 4A). INH treatment brought the bacterial CFU significantly down at both 4 and 8 weeks post-treatment (Fig. 4A). Animals, which received both celecoxib and INH showed more significant reduction in bacterial CFU in the lungs at both 4 and 8 weeks post-treatment (Fig. 4A). Similar results were also obtained in the spleen (Fig. 4B). The combination treatment was significantly more effective with respect to INH or celecoxib alone in controlling bacterial load in spleen at 12 weeks (Fig. 4B).

**Figure 4:**
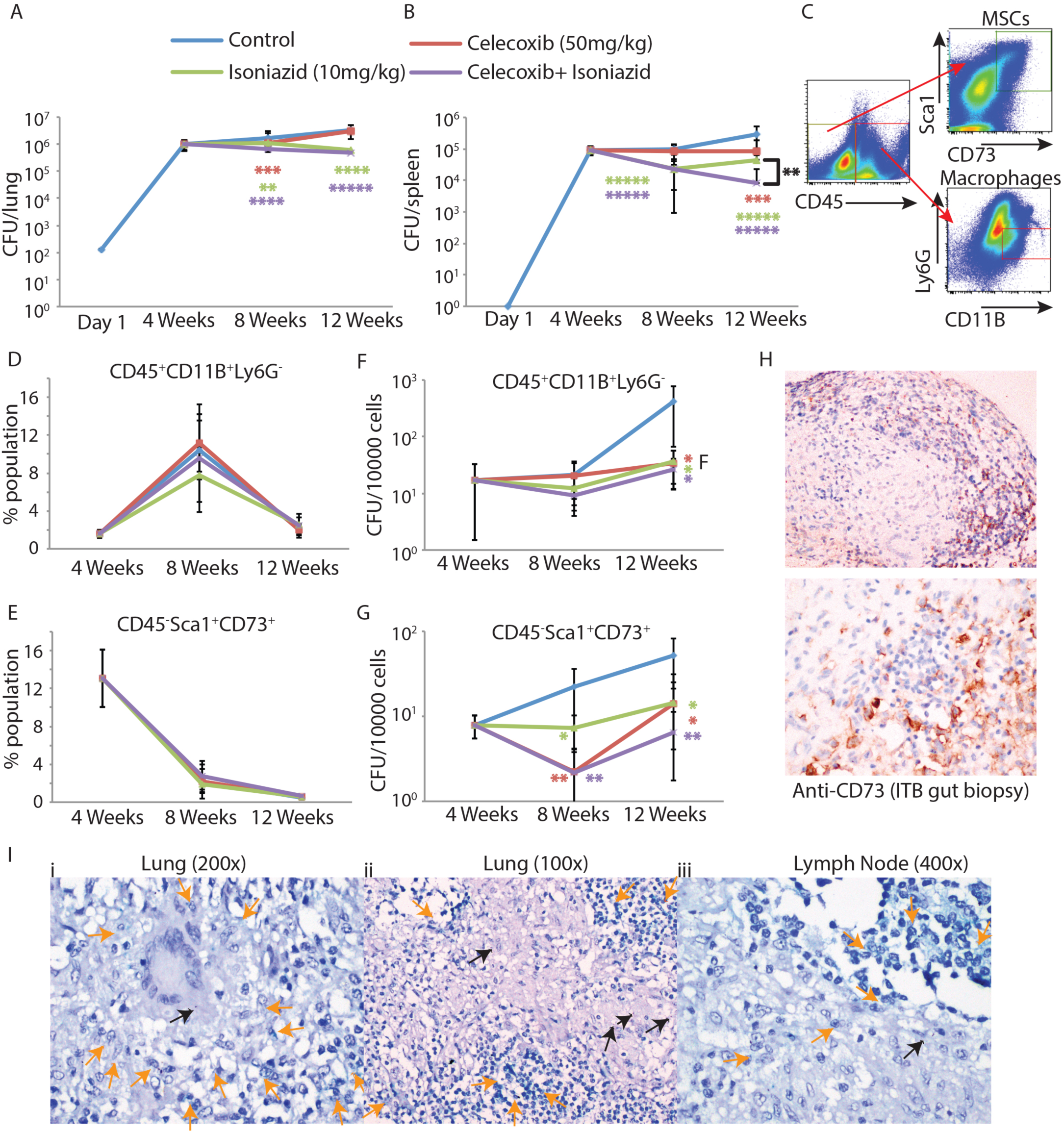
PGE2 facilitates *Mtb* survival within MSCs *in vivo*. **(A)** Total bacterial CFU in the lungs of C57BL/6 mice infected with H37Rv via aerosol route (∼10^2^ bacilli/lung) administered with vector, celecoxib (50 mg/kg), INH (10 mg/kg) or combination of celecoxib with INH (50 mg/kg and 10 mg/kg respectively). Treatments started 4 weeks post-infection and were given every day for next 8 weeks. **(B)** Total bacterial CFU from the spleen of infected animals during the course of experiment mentioned above **(C)** Gating strategy for sorting of MSCs and monocyte/macrophages from mice lung. Live singlet population was gated for CD45 positive and negative population which were sub-gated based on Ly6G^-^CD11b^high^ for macrophages or myeloid cells and Sca1^+^CD73^+^ for MSCs respectively. **(D-E)** Change in tissue landscape with respect to macrophage (CD45^+^Ly6G^-^CD11b^high^) and MSCs across 12 weeks of infection and treatment with celecoxib, INH or celecoxib+INH. **(F-G)** *Mtb* survival within sorted macrophages (CD45^+^Ly6G^-^CD11b^high^) and MSCs (CD45^-^sca1^+^CD73^+^) along the course of infection and treatment as discussed above. **(H)** CD73 staining of biopsies from granuloma positive intestinal tuberculosis patient, showing CD73 positive cells around the submucosal macrogranulomas (x 100, x 200). **(I) (i)** A lung biopsy with an epithelioid cell granuloma surrounded by CD73 positive (orange arrows) [Stay green] cells [x200]. Reddish-brown [AEC positive] *Mtb* bacilli seen within the giant cells and within epithelioid histiocytes (Black arrows). **(ii)** Another lung biopsy shows confluent epithelioid cell granulomas with a few AEC stained bacilli (black arrows). **(iii)** A lymph node biopsy shows *Mtb* antigen positive bacilli (Black arrow), with surrounding CD73 positive cells (Orange arrows) [x 400].

We next sorted lung tissues from the infected animals at each time points and treatment groups into CD45^-^CD73^+^Sca1^+^ (MSCs) and CD45^+^CD11B^+^Ly6G^-^ (macrophages) cells with a purity of more than 90% (Fig. 4C and S5G). Population of macrophages increased dramatically from week 4 to week 8, across all treatment groups and declined subsequently at 12 weeks (Fig. 4D). Population of MSCs was highest at 4 weeks post-infection, which declined rapidly at 8 and 12 weeks post-infection (Fig. 2E). Lysing and plating these sorted cells on 7H11 media showed presence of *Mtb* in both macrophages and MSCs (Fig. 4F-G). Number of bacilli in both cells progressively increased from week 4 to week 12, shown as number of bacilli per ten thousand cells (Fig. 4F and 4G). The macrophage-resident *Mtb* were not affected by celecoxib, INH or both at 4 weeks post-treatment, largely due to low dose of INH used in this study. Moreover, *in vivo* macrophages are also expected to have activated phenotype, thereby could develop drug tolerance as reported previously ^35^. At 8 weeks post-treatment, each of these treatments resulted in significant decline in macrophage-resident *Mtb* population (Fig. 4F). Interestingly, MSC-resident *Mtb* could be killed by INH or celecoxib alone or in combination even at 4 weeks post-treatment (Fig. 4G). However, consistent with the *ex vivo* results, effect of INH alone on MSC-resident *Mtb* was relatively less in magnitude with respect to celecoxib alone or celecoxib+INH treated animals (Fig. 4G). At 8 weeks of treatment, INH and celecoxib were equally effective on MSC-resident *Mtb*, while the combined treatment resulted in additive effect and showed further decline in *Mtb* CFU (Fig. 4G).

### MSCs are present in human pulmonary and extra-pulmonary tuberculosis granulomas

The results so far establish that MSCs serve as a niche for *Mtb* providing drug and immune privileged niche, in PGE2 dependent manner both *ex vivo* and in *vivo in* animals. We next analyzed presence and spatial localization of CD73^+^ cells with respect to *Mtb* in tissue sections from pulmonary and extra-pulmonary TB lesions from human subjects. CD73^+^ cells were found across different pulmonary and extra-pulmonary tuberculosis lesions like gut and lymph nodes (Fig. 4H-I). Intestinal biopsy samples were taken from granuloma-positive confirmed intestinal tuberculosis patients ^49^. In lung and lymph node biopsies where in addition to CD73, Ag85B of *Mtb* was also stained, presence of these cells in the close vicinity of *Mtb* was apparent (Fig. 4I).

## Discussions

Almost everything that we know about the intracellular lifestyle of *Mtb* largely emerged through studies on monocyte/macrophage models. The host responses and mechanism of immune evasions are also studied keeping in mind macrophages as the primary cells where the bacteria reside ^12,13^. The present study was undertaken to understand how mesenchymal stem cells (MSCs) could facilitate mycobacterial persistence in the host as reported by others ^20^. This required us to explore the intracellular lifestyle of *Mtb* within MSCs, not much is known about it. The immune-modulatory properties of MSCs are well known including during *Mtb* infection ^19^. However, in majority of cases, the immune-modulatory effects of MSCs are studied in trans i.e. on a different cell type, which is mediated by effectors released from MSCs ^40-42^. Whether the innate ability of MSCs play a role in mycobacterial persistence and if these cells exhibit any cell-autonomous immune-modulatory properties, is not known. Interestingly, only virulent strain H37Rv could survive and divide well within ADSCs while BCG got killed, suggesting presence of active innate defense mechanism in these cells. One critical aspect of mycobacterial persistence is tolerance to anti-TB drugs, which is driven by host environment like macrophage residence, macrophage activation, low oxygen within granulomas, NO etc. ^35,50^. Our finding that MSC-resident *Mtb* was tolerant to anti-TB drugs underscores the physiological advantage that these cells possess in order to harbor persistent infection as reported previously ^20^. Since adult stem cells are known to have high efflux activity via ABC transporters, which helps in drug tolerance in cases like cancer ^51,52^ we questioned whether these efflux proteins could also help throwing out anti-TB drugs, thereby helping in drug-tolerance. Our results indeed show increased expression and involvement of ABCC1 and ABCG2, in drug tolerance, both of which acted independently since their combined effects were greater than individual effects. However, inhibition of vacuolar-type H^+^ATPase by BafA1 led to a more dramatic decline in drug-tolerant population, suggesting phago-lysosomal environment to be the key factor behind drug-tolerance. Moreover, increase in bacterial CFU upon ABCC1 or ABCG2 inhibition/knockdown in the absence of INH indicated function of these proteins other than cellular efflux of drugs. Their recruitment to bacterial phagosomes indeed point to such a possibility especially since their recruitment to the phagosomes largely correlated with bacterial killing. ABC proteins are known to have several moonlighting functions including nuclear translocation, redox balance and antigen presentation ^53-56^. Results suggest, at least in MSCs, they are also involved in bacterial killing in the phago-lysosomal system. Whether it is associated with lysosomal acidification or transport of bactericidal effectors remains to be uncovered. It is however possible that ABC proteins are actively excluded from getting recruited to *Mtb* phagosomes while being present on other endo-lysosomal vesicles, where through their inwardly transport activities sequester certain anti-bacterial effectors including anti-TB drugs, away from mycobacterial phagosomes in isolated vesicles. This could potentially explain why knocking down or inhibition of ABCC1 or ABCG2 helps increased bacterial survival. At present we have no evidence to support whether these effectors could be H^+^ ions, oxidized glutathione or glutathione metal adducts, ubiquitin-derived peptides and other anti-microbial peptides; each of which are capable of killing the bacteria and are also known targets of ABC proteins-mediated transport across biological membrane ^53,57^.

Mycobacterial drug tolerance can also be induced *in vitro* or *ex vivo* in macrophages. *In vitro, Mtb* develops drug tolerance under stress conditions like hypoxia, NO, nutrient stress etc. ^7,58^. There are also reports, which suggest mere macrophage residence for few hours is sufficient to induce drug tolerance in *Mtb* ^50^. Yet another study reported increased bacterial drug tolerance in activated macrophages ^35^. The common thread across these studies is that when *Mtb* witnesses stress whether *in vitro* or *in vivo*, it activates a set of genes which inadvertently also helps them tide-over the effect of drugs ^11,35^. In agreement with the activated macrophage studies, we noted further increase in drug tolerance of MSC-resident *Mtb* when activated by inflammatory cytokines like IFNγ and TNFα. Most surprisingly though, in the absence of drugs, IFNγ or TNFα treatment did not kill the bacteria, rather helped them grow better. To our knowledge, this is the first report, where under any circumstance a pro-bacterial role for these classic pro-inflammatory cytokines is reported. However MSCs are known to show enhanced immune-modulatory properties when activated with inflammatory cytokines like IFNγ, TNFα and even IL1β ^59,60^. Some key anti-bacterial phenotypes activated in macrophages upon IFNγ stimulation include increased cellular ROS production, mitochondrial depolarization, autophagy inhibition etc. ^39,61^. In MSCs, IFNγ treatment had no such effect and there was significant increase in autophagy. Interestingly, unlike in macrophages, *Mtb* present in MSCs are not present in autophagosomes and therefore xenophagy flux is completely absent ^27,38^. This partially explains the loss of anti-bacterial effects of IFNγ in ADSCs, although does not explain increased bacterial survival upon inflammatory stimuli. However similar to what we noted about ABCC1 or ABCG2 inhibition; pro-bacterial effects of IFNγ and TNFα had mostly to do with lysosomal killing. Since each of these treatments were for the final 24 hours before CFU plating were done, it cannot reflect increased bacterial replication rather show diminished bacterial killing. This observation however brings an exceptionally worrisome insight on the problem of poor efficacy of every vaccine candidates tested so far. While there are several vaccine candidates at different stages of development against tuberculosis to replace or enhance BCG, the only commercially available vaccine, a closer look at each of the vaccine candidate shows that immunological parameters considered as the correlates of protection is common across them^62^. Thus, whether it is MTBVAC or TB/FLU-04L, Ad5Ag85A, MVA85A or others, they all rely on generating strong INFγ producing CD4^+^ and/or CD8^+^ T cells ^63-65^. However given the unconventional pro-bacterial effects of IFNγ on MSCs, these vaccines can only generate an immune response that kills bacterial population in macrophages but not in MSCs thereby blunting the efficacy.

How *Mtb* enjoys such privileged lifestyle within ADSCs became finally apparent through the microarray analysis revealing massive increase in the synthesis and secretion of PGE2 by infected ADSCs. PGE2 is a multifunctional effector, with diverse roles in immune-regulation ^66^. PGE2 mediated immunomodulation of other cells by MSCs has also been extensively reported ^45,46^. However here we report a unique autocrine immune-modulatory function of PGE2 in MSCs. Inhibiting PGE2 signaling was able to revert the pro-bacterial effects of IFNγ, TNFα or MK571, suggesting PGE2 as the converging factor, which helps better bacterial survival within ADSCs. In contrast to the pro-bacterial role of PGE2 observed by us, several studies in the past report protective role of PGE2 against *Mtb*. Thus, loss of PTGES2 makes animals more susceptible to tuberculosis ^67,68^. Similarly, EP2 receptor knockout mice also show increased bacterial burden in the lungs ^69^. Interestingly, PGE2 treatment is more effective in controlling lung CFU only in hyper-susceptible animals lacking IL1R1^67^, with absolutely no effect in WT animals. On the similar line, WT animals and *ptgs2* animals (lacking the enzymatic activity) did not have any difference in bacterial survival ^67^. On the other hand, during late phase of mycobacterial infection and not during early phase of infection, COX2 inhibition has protective effects *in vivo* ^70^. PGE2 is also known to inhibit anti-bacterial effector functions of phagocytes including phagocytosis, NO production, lysosomal killing and antigen presentation ^69^. Incidentally aspirin is currently in clinical trial as adjunct therapy against tuberculosis meningitis in adults ^71^. COX2 inhibitors specially non-steroid anti-inflammatory drugs (NSAIDs) like aspirin, ibuprofen, rofecoxib, celecoxib etc. are routinely used for controlling diverse inflammatory states ^72^. Results from our experiments suggest, smart inclusion of COX2 inhibitors in standard tuberculosis treatment/prevention regimens could enhance the efficacy of treatment. The two major hurdles in the tuberculosis control program are a) lack of effective vaccine and b) highly diminished efficacy of anti-TB drugs *in vivo* with respect to *in vitro*. This study shows MSCs contribute to both these crucial aspects of tuberculosis control. The remodeling of lung granulomas during tuberculosis has been explored previously ^73^. However we show that recruitment and infection of MSCs in the granulomas could be critical events during remodeling considering the lifestyle of *Mtb* in MSCs is radically different than those in macrophages. The study therefore also highlights the limitations of reliance on *ex vivo* data generated through macrophage infection experiments in the past. We believe targeting the immune-privileged environment of MSCs will help develop alternative strategies to enhance both treatment and vaccine efficacy.

## Acknowledgements

The research in DK’s group is supported by Wellcome-DBT India Alliance Senior fellowship (IA/S/17/1/503071). This work was partially supported by DBT, Govt. of India (BT/PR14730/BRB/10/874/2010) and SERB (EMR/2016/005296). NJ and VS are recipients of SRF from CSIR, India and HK has received SRF from UGC. All the work involving *Mycobacterium tuberculosis* including animal infection experiments were performed at Tuberculosis Aerosol Challenge Facility (TACF), a DBT sponsored national facility hosted at ICGEB campus.

## Materials and Methods

Ethical clearance: Studies on human samples were approved by IEC of AIIMS Ref no. IEC-304/02-06-2017 and ICGEB Ref no. ICGEB/IEC/2017/06-verII. Animal experiments were approved by Institutional Animal Ethics Committee, ICGEB (ICGEB/IAEC/280718/CI-14).

### Reagents, antibodies and plasmids

Phorbol 12-myristate 13-acetate (PMA), bafilomycin A1, rapamycin, 3 MA, chloroquine, PKH, *Mtb* drugs (rifampicin,and pyrazinamide), chemical inhibitors (MK571, novobiocin, celecoxib, DMSO, BSA, MTT (1-(4,5-Dimethylthiazol-2-yl)-3,5-diphenylformazan) and paraformaldehyde were obtained from Sigma Aldrich Co (St Louis, MO, USA). Primary antibodies MAP1LC3B, (Cell Signaling Technology and Novus), GAPDH (Santa Cruz Biotechnology), Rab5, ABCC1 (abcam), Rab7, LAMP-1, ABCG2 (santa cruz), CD271, CD73 (BD Bioscience). All IR conjugated secondary antibodies for immunoblotting were obtained from LI-COR Biosciences (Lincoln, NE, USA) while Alexa fluor conjugated secondary antibodies were procured from invitrogen molecular Probes, Carlsbad, CA, USA. PGE2 elisa kit, isoniazid, propidium iodide (PI) were from cayman chemical, USA. Lysotracker red, JC-1 and cellrox green were from Life Technologies, USA. Human IFN-γ and human TNF-α were purchased from ebiosciences. siRNAs (ABCC1, ABCG2) were from GE DHARMACON. Safranin O, Oil Red O and Alizarin Red S was purchased from SRL chemicals.

### Cell culture

Adipose tissue derived mesenchymal stem cells (ADSC) were purchased from life technology (cat no - R7788115) and maintained in mesenPRO RS media (cat no 12746012) supplemented with growth factors at 37°C, 5% CO2, humidified incubator as per the manufacturer instructions and guidelines. For all *in vitro* experiments, ADSC were seeded at the required density, allowed to adhere to the surface for 24-36 hours before proceeding with the experiment.

Human monocytic cell line THP-1 were obtained from American type culture collection (ATCC) and cultured in RPMI 1640 medium along with 10% fetal bovine serum (FBS) at 37°C, 5% CO_2_ humidified incubator. THP-1 derived macrophages (dTHP-1) were obtained by treating THP-1 cells with 20ng/ml phorbol myristate acetate (PMA, sigma) for 24 hours followed by PMA removal and maintenance in complete media.

### MSC Characterization

MSC characterization into 3 lineages i.e. osteocytes, chondrocytes and adipocytes were done according to the differentiation protocol by life technologies. In brief, MSC were seeded in 24 well plate and after 6-8 hours of adherence their media was replaced with adipocyte differentiation media (cat no A1007001), chondrocyte differentiation media (cat no A1007101) and osteocyte differentiation media (cat no A1007201). Media was replaced every 3^rd^ day till 14^th^ day. After 14 days, cells were fixed and stained with Alizarin red S for chondrocytes, Oil Red O for adipocytes, Safranin O for osteocytes.

### Bacterial cultures and *in vitro* infection experiments

Virulent laboratory strain H37Rv and vaccine strain BCG bacterial cultures were grown in 7H9 media (BD Difco) supplemented with 10% Albumin - Dextrose - Catalase (ADC, BD Difco) and incubated in an orbitary shaker at 100 rpm, 37° C until the mid-log phase. Single cell suspension required for carrying out infection experiments were prepared by passing bacterial culture through a series of different gauge needles, five times through 23 gauge and 26 gauge and thrice through 30 gauge.

For macrophage experiments, bacterial infection was set up for 4 hours followed by RPMI wash and addition of amikacin sulphate at a final concentration of 200µg/ml for 2 hours to kill any extracellular bacteria. For ADSCs, infection was done for 12 hours followed by addition of amikacin sulphate (200µg/ml) for 2 hours and replenishment of fresh media. All the treatments of drug, cytokines, autophagy modulators or chemical inhibitors were done 24 hours before the 6^th^ day and 3^rd^ day time point for ADSC and dTHP-1 respectively while siRNA transfections were done 48hrs before the 6^th^ day and 3^rd^ day time point for ADSC and dTHP-1 respectively. Bacterial colony forming units (CFU) were enumerated by adding lysis buffer, 7H9 containing 0.1% SDS in the required plate and incubating for 5 min and plating on 7H11 agar plates supplemented with OADC (BD Difco). The plates were incubated at 37°C to allow bacterial growth, and counts were performed after 21 days. Percent tolerant population was calculated for CFU obtained in drug treated set as percent of the CFU in the untreated set from the same experiment.

### MTT Assay

Cell viability was assessed by performing MTT (sigma) [3-(4,5-dimethyl-2-thiazolyl)-2,5-diphenyl-2H-tetrazolium bromide] assay. At indicated time points, media was removed from the plate and washed once with phenol free RMPI. MTT was prepared in phenol free RPMI at a working concentration of 1 mg/ml. 100 µl of MTT solution was added to each well of 96 well plates and incubated for appropriate time in cell incubator. Thereafter, MTT solution was removed and formazan crystals were dissolved in 100 µl DMSO. Samples obtained thereafter were quantified by measuring their absorbance at 560 nm in the plate reader.

### Confocal microscopy

For confocal experiments, bacteria were stained with CellVue@claret dye (sigma) or PKF67, far-red/green lipophilic dye, according to the manufactures protocol and resuspended into final media and incubated with cells for infection. To visualize acidified compartments, LysoTracker red dye (LysoTracker Red DND-99; Life Technologies) was added to the sample wells at a concentration of 500 nM for 30 mins. Cells were fixed in 4% (w/v) PFA for 15-20 mins, followed by PBS wash twice and ammonium chloride treatment for 15 mins. Cells were again washed with PBS and incubated with 0.2 % TritonX-100 in 1X PBS for 20 min to ensure permeabilization. It was followed by blocking solution (3% BSA in 1X PBST) for 1 hour. Cells were then incubated with primary antibody for 2-3 hours at RT, followed by PBST wash and conjugated secondary antibody for an hour. Cells were given a final wash with 1X PBS and coverslips were mounted in ProLong Gold antifade reagent (Life Technologies). Images were acquired by NIS-Elements software using the Nikon A1R laser scanning confocal microscope equipped with a Nikon Plan Apo 60× 1.40-numerical-aperture (NA) oil immersion objective. Serial confocal sections, 0.5 µm thick, were acquired within a *z*-stack spanning 10 to 15 µm to form a composite image. Images were analysed using Imaris and image J software

### Flow cytometry

Surface and intracellular protein expression in THP-1 and ADSC were carried out using flow cytometry. At required time points, cells were scrapped off, pelleted and washed. Cells were pelleted down at 1000 rpm and blocked in 3% BSA in 1X PBS and incubated with primary antibody for 3 hr in blocking buffer followed by incubation with alexa flour 488 conjugated secondary antibody for 2 hrs (surface expression). Intracellular expression was assessed after permeabilizing cells with 0.05% saponin, followed by blocking, primary and secondary antibody incubation. After incubations cells were washed with 1X PBS and re-suspended in 1X PBS and samples were acquired in BD FACS Canto II by using FACS Diva acquisition software. For measurement of cellular ROS, cells were scrapped at required time point and stained with CellROX Green before acquisition on BD FACS Canto II. Staining of the dyes were performed as per the manufacturer’s directions. The data was analyzed using Flow Jo V. software.

### Real time PCR and Microarray

Total RNA from ADSC and dTHP-1 cells was isolated using mdi RNA isolation kit. cDNA was synthesized from 500ng of total RNA by reverse transcriptase PCR using Bio-RAD iScript cDNA synthesis kit according to the manufacturer’s protocol. ADSC and dTHP-1 cDNA samples was run in triplicate using β-tubulin and actin as normalizing control respectively using SYBR green dye for real time fluorescence acquisition on the Bio-Rad CFX 96 Real time PCR system. Primers were synthesized from Sigma Aldrich Chemicals Ltd. Primers used: ABCC1 (F:CGAGAACCAGAAGGCCTATTAC, R:ACAGGGCAGCAAACAGAA) ABCG2 (F:CTTCGGCTTGCAACAACTATG,R:CCAGACACACCACGGATAAA), Tubulin (F:TTGGCCAGATCTTTAGACCAGACAAC,R: CCGTACCACATCCAGGACAGAATC),Actin (F: ACCTTCTACAATGAGCTGCG, R: CCTGGATAGCAACGTACATGG)

For microarray, total RNA from ADSC uninfected or infected H37Rv was extracted using MDI RNA isolation kit. Samples were sent to Bionivid Technologies, Bangalore for cDNA synthesis and hybridization to 25 “Illumina human WholeGenome-6 version 2 BeadChips” using standard illumina protocols. Six biological replicates were used for hybridization.

### C57BL/6 aerosol challenge

All mice experiments were carried out in the Tuberculosis Aerosol Challenge Facility (TACF, ICGEB, New Delhi, India). C56BL/6 mice were housed in cages contained within a biosafety level 3 laminar flow enclosure. Aerosol challenge of 100 CFU was given to the animals in a Wisconsin-Madison chamber according to the standardized protocol. To check for infection establishment, two animals were selected randomly and humanely euthanized 24 hours post-aerosol challenge. The lungs and spleen tissues were harvested and homogenized to enumerate CFU. Tissue lysates were serially diluted and plated on petri plates containing Middlebrook 7H11 agar (Difco) supplemented with 10% OADC (Becton, Dickinson) and 0.5% glycerol.

### Animal dosing, CFU plating and FACS sorting

For *in vivo* experiments, drug dosing was initiated 4 weeks post-aerosol challenge and the animals were administered the drug by oral gavage at a dose of 10 mg/kg INH and 50 mg/kg celecoxib in combination and individually every day till 12 weeks. After 8 and 12 weeks, 6 mice in each group were euthanized, followed by removal of lung and spleen and their homogenization and plating. Half lung was used in the preparation of single cell suspension followed by FACS sorting and subsequent plating. To begin with the tissue was washed with PBS, chopping it in small pieces followed by addition of 20U/ml DNAses, 1mg/ml collagenase D and incubation for 30 mins at 37C. This is passed through nylon mesh to get single cell suspension. The cells were pelleted at 2000 rpm, treated with RBC lysis buffer, washed with PBS. Cells were stained with the antibody cocktail of CD45, CD73, CD11b, ly6G, sca-1 and just before sorting PI were added at 5µg/ml for live/dead staining. All the cells were sorted to get the maximum number of sorted cells which were then pelleted, lysed and plated.

### Immunohistochemistry

Five-micron thick sections of tissue were taken on the coated slide. Deparaffinization was done by dipping the slides in xylene for 5 mins (2 changes), acetone for 2-3 mins, alcohol for 2-3 mins and then under running/tap water. Antigen retrieval was performed with citrate buffer (pH=6) in microwave oven, at 100°C for 30 minutes. Endogenous persxidase blocking was done with 4% Hydrogen peroxide in methanol. Rabbit anti-Ag85B primary antibody was added (1:400) and incubated overnight. Thereafter goat anti-rabbit IgG H&L (ab6721) was added and incubated for 40 minutes followed by 200 µl of enhancer for 5 minutes at RT. For color development, 3-amino-9-ethiylcarbazole chromogen was used at RT for 15 minutes. Thereafter the Mouse anti-CD73 antibody (ab91086, abcam, Cambridge, USA) was added in a dilution of 1:500 overnight at RT. Alkaline phosphatase tagged goat anti-mouse IgG H&L secondary antibody (ab7069) was added (1:200) for 40 minutes at room temperature. Color was developed using a Stay Green/AP plus kit (ab156428) following manufacturer’s protocol.

### Statistical analysis

Statistical significance for comparisons between two sets of the experiments was done using unpaired two-tailed Students’s *t*-test. **denotes significant difference at *P*<0.01 and * at *P*<0.05.

## References

1. Russell, D. G., Barry, C. E., 3rd & Flynn, J. L. Tuberculosis: what we don’t know can, and does, hurt us. Science (New York, N.Y.) 328, 852N856, doi:10.1126/science.1184784 (2010).

2. Voss, G. et al. Progress and challenges in TB vaccine development. F1000Research 7, 199N199, doi:10.12688/f1000research.13588.1 (2018).

3. Streicher, E. M. et al. Emergence and treatment of multidrug resistant (MDR) and extensively drugNresistant (XDR) tuberculosis in South Africa. Infection, Genetics and Evolution 12, 686N694, doi:https://doi.org/10.1016/j.meegid.2011.07.019 (2012).

4. Ehlers, S. & Schaible, U. E. The granuloma in tuberculosis: dynamics of a hostNpathogen collusion. Frontiers in immunology 3, 411N411, doi:10.3389/fimmu.2012.00411 (2013).

5. Boshoff, H. I. & Barry, C. E., 3rd. Tuberculosis N metabolism and respiration in the absence of growth. Nature reviews. Microbiology 3, 70N 80, doi:10.1038/nrmicro1065 (2005).

6. CunninghamNBussel, A., Zhang, T. & Nathan, C. F. Nitrite produced by Mycobacterium tuberculosis in human macrophages in physiologic oxygen impacts bacterial ATP consumption and gene expression. Proceedings of the National Academy of Sciences of the United States of America 110, E4256NE4265, doi:10.1073/pnas.1316894110 (2013).

7. Wayne, L. G. & Hayes, L. G. An in vitro model for sequential study of shiftdown of Mycobacterium tuberculosis through two stages of nonreplicating persistence. Infect Immun 64, 2062N2069 (1996).

8. Yang, C.NS., Yuk, J.NM. & Jo, E.NK. The role of nitric oxide in mycobacterial infections. Immune network 9, 46N52, doi:10.4110/in.2009.9.2.46 (2009).

9. Flentie, K., Garner, A. L. & Stallings, C. L. Mycobacterium tuberculosis Transcription Machinery: Ready To Respond to Host Attacks. Journal of bacteriology 198, 1360N1373, doi:10.1128/JB.00935N15 (2016).

10. Mehta, M., Rajmani, R. S. & Singh, A. Mycobacterium tuberculosis WhiB3 Responds to Vacuolar pHNinduced Changes in Mycothiol Redox Potential to Modulate Phagosomal Maturation and Virulence. The Journal of biological chemistry 291, 2888N2903, doi:10.1074/jbc.M115.684597 (2016).

11. Mehta, M. & Singh, A. Mycobacterium tuberculosis WhiB3 maintains redox homeostasis and survival in response to reactive oxygen and nitrogen species. Free Radic Biol Med 131, 50N58, doi:10.1016/j.freeradbiomed.2018.11.032 (2018).

12. Cosma, C. L., Sherman, D. R. & Ramakrishnan, L. The secret lives of the pathogenic mycobacteria. Annu Rev Microbiol 57, 641N676, doi:10.1146/annurev.micro.57.030502.091033 (2003).

13. Guirado, E., Schlesinger, L. S. & Kaplan, G. Macrophages in tuberculosis: friend or foe. Seminars in immunopathology 35, 563N583, doi:10.1007/s00281N013N0388N2 (2013).

14. BeigierNBompadre, M. et al. Mycobacterium tuberculosis infection modulates adipose tissue biology. PLOS Pathogens 13, e1006676, doi:10.1371/journal.ppat.1006676 (2017).

15. Lyadova, I. V. Neutrophils in Tuberculosis: Heterogeneity Shapes the Way? Mediators of Inflammation 2017, 8619307, doi:10.1155/2017/8619307 (2017).

16. Russell, D. G., Cardona, P.NJ., Kim, M.NJ., Allain, S. & Altare, F. Foamy macrophages and the progression of the human TB granuloma. Nature immunology 10, 943N948, doi:10.1038/ni.1781 (2009).

17. Scordo, J. M., Knoell, D. L. & Torrelles, J. B. Alveolar Epithelial Cells in Mycobacterium tuberculosis Infection: Active Players or Innocent Bystanders? Journal of innate immunity 8, 3N14, doi:10.1159/000439275 (2016).

18. Khan, A. et al. Mesenchymal stem cells internalize Mycobacterium tuberculosis through scavenger receptors and restrict bacterial growth through autophagy. Scientific Reports 7, 15010, doi:10.1038/s41598N017N 15290Nz (2017).

19. Raghuvanshi, S., Sharma, P., Singh, S., Van Kaer, L. & Das, G. Mycobacterium tuberculosis evades host immunity by recruiting mesenchymal stem cells. Proceedings of the National Academy of Sciences of the United States of America 107, 21653N21658, doi:10.1073/pnas.1007967107 (2010).

20. Das, B. et al. CD271(+) Bone Marrow Mesenchymal Stem Cells May Provide a Niche for Dormant Mycobacterium tuberculosis. Science translational medicine 5, 170ra113N170ra113, doi:10.1126/scitranslmed.3004912 (2013).

21. Zhang, Y., Yew, W. W. & Barer, M. R. Targeting persisters for tuberculosis control. Antimicrobial agents and chemotherapy 56, 2223N2230, doi:10.1128/AAC.06288N11 (2012).

22. Gomez, J. E. & McKinney, J. D. M. tuberculosis persistence, latency, and drug tolerance. Tuberculosis (Edinburgh, Scotland) 84, 29N44 (2004).

23. Wakamoto, Y. et al. Dynamic persistence of antibioticNstressed mycobacteria. Science (New York, N.Y.) 339, 91N95, doi:10.1126/science.1229858 (2013).

24. Bernardo, M. E. & Fibbe, W. E. Mesenchymal stromal cells: sensors and switchers of inflammation. Cell Stem Cell 13, 392N402, doi:10.1016/j.stem.2013.09.006 (2013).

25. Singer, N. G. & Caplan, A. I. Mesenchymal stem cells: mechanisms of inflammation. Annu Rev Pathol 6, 457N478, doi:10.1146/annurevNpatholN 011110N130230 (2011).

26. Zhao, Q., Ren, H. & Han, Z. Mesenchymal stem cells: Immunomodulatory capability and clinical potential in immune diseases. Journal of Cellular Immunotherapy 2, 3N20, doi:https://doi.org/10.1016/j.jocit.2014.12.001 (2016).

27. Chandra, P. et al. Mycobacterium tuberculosis Inhibits RAB7 Recruitment to Selectively Modulate Autophagy Flux in Macrophages. Scientific Reports 5, 16320, doi:10.1038/srep16320 https://www.nature.com/articles/srep16320 - supplementary-Ninformation (2015).

28. Karim, A. F. et al. Express Path Analysis Identifies a Tyrosine Kinase SrcN centric Network Regulating Divergent Host Responses to Mycobacterium tuberculosis Infection. The Journal of biological chemistry 286, 40307N 40319, doi:10.1074/jbc.M111.266239 (2011).

29. Kumar, D. et al. GenomeNwide Analysis of the Host Intracellular Network that Regulates Survival of Mycobacterium tuberculosis. Cell 140, 731N743, doi:https://doi.org/10.1016/j.cell.2010.02.012 (2010).

30. Gottesman, M. M., Fojo, T. & Bates, S. E. Multidrug resistance in cancer: role of ATPNdependent transporters. Nat Rev Cancer 2, 48N58, doi:10.1038/nrc706 (2002).

31. Szakacs, G., Paterson, J. K., Ludwig, J. A., BoothNGenthe, C. & Gottesman, M. M. Targeting multidrug resistance in cancer. Nat Rev Drug Discov 5, 219N 234, doi:10.1038/nrd1984 (2006).

32. Zaman, G. J. et al. The human multidrug resistanceNassociated protein MRP is a plasma membrane drugNefflux pump. Proceedings of the National Academy of Sciences of the United States of America 91, 8822N8826 (1994).

33. Lemos, C. et al. Folate deprivation induces BCRP (ABCG2) expression and mitoxantrone resistance in Caco-2 cells. International Journal of Cancer 123, 1712N1720, doi:10.1002/ijc.23677 (2008).

34. Klionsky, D. et al. Guidelines for the use and interpretation of assays for monitoring autophagy (3rd edition). Vol. 12 (2016).

35. Liu, Y. et al. Immune activation of the host cell induces drug tolerance in Mycobacterium tuberculosis both in vitro and in vivo. The Journal of experimental medicine 213, 809N825, doi:10.1084/jem.20151248 (2016).

36. MacGurn, J. A. & Cox, J. S. A Genetic Screen for ≪em≫Mycobacterium tuberculosis≪/em≫ Mutants Defective for Phagosome Maturation Arrest Identifies Components of the ESXN1 Secretion System. Infection and Immunity 75, 2668, doi:10.1128/IAI.01872N06 (2007).

37. Deretic, V. Autophagy, an immunologic magic bullet: Mycobacterium tuberculosis phagosome maturation block and how to bypass it. Future microbiology 3, 517N524, doi:10.2217/17460913.3.5.517 (2008).

38. Chandra, P. & Kumar, D. Selective autophagy gets more selective: Uncoupling of autophagy flux and xenophagy flux in Mycobacterium tuberculosisNinfected macrophages. Autophagy 12, 608N609, doi:10.1080/15548627.2016.1139263 (2016).

39. Matta, S. K. & Kumar, D. AKT mediated glycolytic shift regulates autophagy in classically activated macrophages. The International Journal of Biochemistry & Cell Biology 66, 121N133, doi:https://doi.org/10.1016/j.biocel.2015.07.010 (2015).

40. Ghannam, S., Bouffi, C., Djouad, F., Jorgensen, C. & Noel, D. Immunosuppression by mesenchymal stem cells: mechanisms and clinical applications. Stem cell research & therapy 1, 2, doi:10.1186/scrt2 (2010).

41. Nemeth, K. et al. Bone marrow stromal cells attenuate sepsis via prostaglandin E(2)Ndependent reprogramming of host macrophages to increase their interleukinN10 production. Nature medicine 15, 42N49, doi:10.1038/nm.1905 (2009).

42. Najar, M. et al. Mesenchymal stromal cells use PGE2 to modulate activation and proliferation of lymphocyte subsets: Combined comparison of adipose tissue, Wharton’s Jelly and bone marrow sources. Cellular Immunology 264, 171N179, doi:https://doi.org/10.1016/j.cellimm.2010.06.006 (2010).

43. Corcione, A. et al. Human mesenchymal stem cells modulate BNcell functions. Blood 107, 367N372, doi:10.1182/bloodN2005N07N2657 (2006).

44. Ling, W. et al. Mesenchymal stem cells use IDO to regulate immunity in tumor microenvironment. Cancer research 74, 1576N1587, doi:10.1158/0008N5472.canN13N1656 (2014).

45. Wang, B. et al. mTOR inhibition improves the immunomodulatory properties of human bone marrow mesenchymal stem cells by inducing COXN2 and PGE2. Stem cell research & therapy 8, 292, doi:10.1186/s13287N017N0744N6 (2017).

46. Zhang, W. et al. Effects of mesenchymal stem cells on differentiation, maturation, and function of human monocyteNderived dendritic cells. Stem cells and development 13, 263N271, doi:10.1089/154732804323099190 (2004).

47. Kalle, A. M. & Rizvi, A. Inhibition of bacterial multidrug resistance by celecoxib, a cyclooxygenaseN2 inhibitor. Antimicrob Agents Chemother 55, 439N442, doi:10.1128/AAC.00735N10 (2011).

48. Reid, G. et al. The human multidrug resistance protein MRP4 functions as a prostaglandin efflux transporter and is inhibited by nonsteroidal antiinflammatory drugs. Proceedings of the National Academy of Sciences of the United States of America 100, 9244N9249, doi:10.1073/pnas.1033060100 (2003).

49. Sharma, R., Madhusudhan, K. S. & Ahuja, V. Intestinal tuberculosis versus crohn’s disease: Clinical and radiological recommendations. The Indian journal of radiology & imaging 26, 161N172, doi:10.4103/0971N 3026.184417 (2016).

50. Adams, K. N. et al. Drug tolerance in replicating mycobacteria mediated by a macrophageNinduced efflux mechanism. Cell 145, 39N53, doi:10.1016/j.cell.2011.02.022 (2011).

51. Sharom, F. J. ABC multidrug transporters: structure, function and role in chemoresistance. Pharmacogenomics 9, 105N127, doi:10.2217/14622416.9.1.105 (2007).

52. Singh, A. et al. Expression of ABCG2 (BCRP), a Marker of Stem Cells, is Regulated by Nrf2 in Cancer Cells That Confers Side Population and Chemoresistance Phenotype. Molecular cancer therapeutics 9, 2365N2376, doi:10.1158/1535N7163.MCTN10N0108 (2010).

53. Cole, S. P. C. & Deeley, R. G. Transport of glutathione and glutathione conjugates by MRP1. Trends in Pharmacological Sciences 27, 438N446, doi:10.1016/j.tips.2006.06.008 (2006).

54. Liang, S.NC. et al. ABCG2 Localizes to the Nucleus and Modulates CDH1 Expression in Lung Cancer Cells()()(). Neoplasia (New York, N.Y.) 17, 265N 278, doi:10.1016/j.neo.2015.01.004 (2015).

55. Seyffer, F. & Tampe, R. ABC transporters in adaptive immunity. Biochimica et biophysica acta 1850, 449N460, doi:10.1016/j.bbagen.2014.05.022 (2015).

56. Chandra, S. et al. Mrp1 is involved in lipid presentation and iNKT cell activation by Streptococcus pneumoniae. Nat Commun 9, 4279, doi:10.1038/s41467N018N06646N8 (2018).

57. Bryan, J. et al. ABCC8 and ABCC9: ABC transporters that regulate K+ channels. Pflugers Archiv: European journal of physiology 453, 703N718, doi:10.1007/s00424N006N0116Nz (2007).

58. Hampshire, T. et al. Stationary phase gene expression of Mycobacterium tuberculosis following a progressive nutrient depletion: a model for persistent organisms? Tuberculosis (Edinburgh, Scotland) 84, 228N238, doi:10.1016/j.tube.2003.12.010 (2004).

59. Broekman, W. et al. TNFNalpha and ILN1betaNactivated human mesenchymal stromal cells increase airway epithelial wound healing in vitro via activation of the epidermal growth factor receptor. Respir Res 17, 3, doi:10.1186/s12931N015N0316N1 (2016).

60. Ryan, J. M., Barry, F., Murphy, J. M. & Mahon, B. P. InterferonNgamma does not break, but promotes the immunosuppressive capacity of adult human mesenchymal stem cells. Clin Exp Immunol 149, 353N363, doi:10.1111/j.1365N2249.2007.03422.x (2007).

61. Matta, S. K. & Kumar, D. Hypoxia and classical activation limits Mycobacterium tuberculosis survival by AktNdependent glycolytic shift in macrophages. Cell Death Discovery 2, 16022, doi:10.1038/cddiscovery.2016.22 https://www.nature.com/articles/cddiscovery201622 N supplementaryN information (2016).

62. Atmakuri, K., PennNNicholson, A., Tanner, R. & Dockrell, H. M. Meeting report: 5th Global Forum on TB Vaccines, 20N23 February 2018, New Delhi India. Tuberculosis (Edinburgh, Scotland) 113, 55N64, doi:10.1016/j.tube.2018.08.013 (2018).

63. Bertholet, S. et al. Identification of human T cell antigens for the development of vaccines against Mycobacterium tuberculosis. Journal of immunology (Baltimore, Md.: 1950) 181, 7948N7957 (2008).

64. Lewinsohn, D. A., Lewinsohn, D. M. & Scriba, T. J. Polyfunctional CD4(+) T Cells As Targets for Tuberculosis Vaccination. Frontiers in immunology 8, 1262N1262, doi:10.3389/fimmu.2017.01262 (2017).

65. Ritz, N., Hanekom, W. A., RobinsNBrowne, R., Britton, W. J. & Curtis, N. Influence of BCG vaccine strain on the immune response and protection against tuberculosis. FEMS Microbiology Reviews 32, 821N841, doi:10.1111/j.1574N6976.2008.00118.x (2008).

66. Kalinski, P. Regulation of immune responses by prostaglandin E2. Journal of immunology (Baltimore, Md.: 1950) 188, 21N28, doi:10.4049/jimmunol.1101029 (2012).

67. MayerNBarber, K. D. et al. HostNdirected therapy of tuberculosis based on interleukinN1 and type I interferon crosstalk. Nature 511, 99N103, doi:10.1038/nature13489 (2014).

68. Divangahi, M. et al. Mycobacterium tuberculosis evades macrophage defenses by inhibiting plasma membrane repair. Nat Immunol 10, 899N 906, doi:10.1038/ni.1758 (2009).

69. Kaul, V. et al. An important role of prostanoid receptor EP2 in host resistance to Mycobacterium tuberculosis infection in mice. J Infect Dis 206, 1816N1825, doi:10.1093/infdis/jis609 (2012).

70. Rangel Moreno, J. et al. The role of prostaglandin E2 in the immunopathogenesis of experimental pulmonary tuberculosis. Immunology 106, 257N266 (2002).

71. Mai, N. T. et al. A randomised double blind placebo controlled phase 2 trial of adjunctive aspirin for tuberculous meningitis in HIVNuninfected adults. eLife 7, doi:10.7554/eLife.33478 (2018).

72. Crofford, L. J. Use of NSAIDs in treating patients with arthritis. Arthritis research & therapy 15 Suppl 3, S2NS2, doi:10.1186/ar4174 (2013).

73. Dheda, K. et al. Lung remodeling in pulmonary tuberculosis. J Infect Dis 192, 1201N1209, doi:10.1086/444545 (2005).

