## Supplemental Material (Figures and legends) for "Mesenchymal stem cells (MSCs) offer a drug tolerant and immune- privileged niche to *Mycobacterium tuberculosis*"

**Contains legends for:**

**Supplementary figures: S1-S5**

**Supplementary Table: S1**

**Supplemental Figure S1: Infection of ADSCs with H37Rv**

**(A)** Line histogram of ADSCs stained with CD73-FITC and CD-271 PE-cy7. **(B)** Uninfected ADSCs differentiated into osteocytes, chondrocytes and adipocytes using specific differentiation media and stained with safranin O, Alizarin red S and oil red O respectively, visualized under light microscope at 10X (Osteocytes, chondriocytes) and 40X (Adipocytes). **(C)** 6$^{th}$ day H37Rv infected ADSCs, without any differentiation media, stained with safranine O, Alizarin Red S, and oil red O compared to uninfected differentiated ADSCs as positive controls. Images are taken at 10X and 40X respectively using light microscope. **(D)** Volcano plot depicting differentially regulated genes (depicted in blue) in ADSCs upon *Mtb* infection with respect to uninfected ADSCs with p-value at y-axis and $\log_2$ fold change on x-axis. **(E)** Graph highlighting the fold change in genes involved in differentiation and maintenance of chondrocytes, adipocytes and osteocytes as determined from microarray analysis of *Mtb* infected ADSCs 6$^{th}$ day post-infection. **(F)** Circos plot showing the enriched gene ontology pathways (GO) regulated upon infection of ADSCs with H37Rv. Each pathway highlighted with different colour (right half of the circle) is linked to the individual genes (left half of the circle), which are either up or down regulated shown in colors depicted in the $\log_2$ fold change pseudocolor scale.

**Supplemental Figure S2: Role of efflux pumps ABCC1 and ABCG2 in determining bacterial phenotypes within ADSCs**

**(A)** Fold change in mRNA expression of ABCC1 and ABCG2 in ADSCs infected with different MOI of bacteria i.e.1, 5, 10, 25 and 50 on 6$^{th}$ day post-infection as determined by qPCR. β-tubulin was used as a constitutive housekeeping gene and data was normalized to uninfected ADSCs. **(B-C)** Percent INH tolerant H37Rv population in ADSCs after addition of novobiocin (ABCG2 inhibitor, 25 μM) (B) or doses of MK571 (ABCC1 inhibitor), 10, 25, 50 μM (C) for 24 hours before CFU plating on 6$^{th}$ day post-infection. **(D-E)** *In vitro* growth of H37Rv broth culture in the presence of (D) MK571 (10,25,50,75,100 μM) along with vehicle control (DMSO) or novobiocin (10, 25, 50, 75, 100 μg/ml) for 24 hours and measured spectrophotometrically (O.D. 600 nm). **(F)** Line histogram of intracellular (I.C.) and surface expression of ABCC1/MRP-1 and ABCG2/BCRP in uninfected and 6$^{th}$ day H37Rv infected ADSCs with and without siRNA. Red line represents the Isotype secondary control. **(G-H)** CFU assay of *Mtb* load in ADSCs on 6$^{th}$ day post-infection, after 24 hours treatment with different doses of MK571 (F) and novobiocin (G). Data represented is cumulative of 3 independent experiments. Error bar represent S.E.M. *P < 0.05, **P < 0.005, ***P < 0.0005, ****P < 0.00005, NS 'not significant' by two-tailed Student's t-test.

**Supplemental Figure S3: Intracellular niches of H37Rv within ADSCs**

**(A-B)** Representative confocal images of PKH67 labeled H37Rv infected ADSCs stained with Rab5 along with DAPI (A) and Rab7 with DAPI (B) 3$^{rd}$ day post-infection. Bar graphs at the right shows percent localization of *Mtb* with Rab5 and Rab7 compartments respectively. Data is cumulative of 10 fields with >50 bacteria and experiment repeated at least thrice. **(C)** Percent localization of PKH67 labeled H37Rv to Rab5 and Rab7 compartments in THP1 macrophages at 48 hours post-infection **(D)** Confocal images PKH67 labeled H37Rv infected ADSCs stained for lysosomal markers i.e. LAMP-1, Cathepsin D and LysoTracker Red after 24 hour incubation with IFNγ (100 units/ml), TNFα (20 ng/ml) or MK571 (50 μM) or left untreated on 3$^{rd}$ day post-infection. **(E)** PKH67 labeled H37Rv infected ADSCs stained with ABCC1 or ABCG2 along with lysoTracker red. Scale bars, 10 μm.

**Supplemental Figure S4: Autophagy regulation in ADSCs upon *Mtb* infection and immune activation**

**(A)** Representative confocal images of GFP-H37Rv infected ADSCs stained with LC3 and DAPI without or with the addition of BafA1 (100 nM) for 3 hours, 3$^{rd}$ day post-infection. Bar graph at the right represents percent localization of *Mtb* with LC3 compartment on Day 0 and Day 3 (top) and on addition of BafA1 on (bottom). **(B)** Immunoblot of uninfected ADSCs probed for LC3 protein with GAPDH as loading control after addition of rapamycin (100 nM, 6 hours) and starvation/HBSS (6 hours) in the absence or presence of BafA1 (100 nM, 3 hours). Immunoblot for LC3 was also performed for uninfected and 6$^{th}$ day H37Rv infected ADSCs with and without BafA1 (100 nM, 3 hours) with GAPDH as loading control. **(C)** Immunoblot of uninfected and 6$^{th}$ day H37Rv infected ADSCs probed for LC3 protein and GAPDH upon 24 hours treatment with 100 units/ml IFNγ and 20 ng/ml TNF with and without the addition of BafA1 (100 nM, 3 hours). **(D)** Line histogram of cellular ROS measured using DCFDA dye (5 μM, 30 minutes) within uninfected and infected ADSCs left untreated or treated with increasing doses of IFNγ (100, 250 and 500 units/ml). Scale bar, 10 μm. Error bars represent S.E.M.

**Supplemental Figure S5: PGE2 plays key role in providing the protective niche to *Mtb* within ADSCs**

Bar graph depicts the $\log_2$ fold change in the expression of genes known for MSC-mediated immunomodulation, in ADSCs on 6th day post-infection in the microarray analysis (Table S1). **(B)** CFU burden of 6th day infected ADSC when left untreated or treated with IFNγ, TNFα and MK571 along with increasing doses of PF04418948 (50, 250 and 500 nM). **(C)** CFU assay in infected THP-1 macrophages after addition of increasing dose of celecoxib (50, 150, 250 μM) for 24 hours prior to 3rd day plating. **(D)** PGE2 Elisa of supernatnats from uninfected and H37Rv infected THP-1 macrophages collected on 3rd day pot-infection, which were also untreated or treated with IFNγ (100 units/ml), TNFα (20 ng/ml), and celecoxib (250 μM) for 24 hours, done according to the manufacturer's protocol. **(E)** Immunoblot showing knowdown of cox-2 protein 48 hours after transfecting 100 nM and 200 nM of cox-2 siRNA or scrambled control. GAPDH is used as a loading control. **(F)** Representative confocal images of PKH67 labelled H37Rv infected ADSCs dual stained with LAMP-1 and CatD along with LysoTracker red after treatment with celecoxib (250 μM, 24 hours) or PF (500 nM, 24 hours) or cox-2 knockdown (100 nM siRNA, 48 hours) 3rd day post-infection. Scale bars, 10 μm. Error bars represent S.E.M. **(G)** Percent purity of sorted MSCs and macrophages.

**Supplementary Table S1: Table enlisting gene expression profile of ADSCs upon infection with H37Rv**

Supplementary figure S1

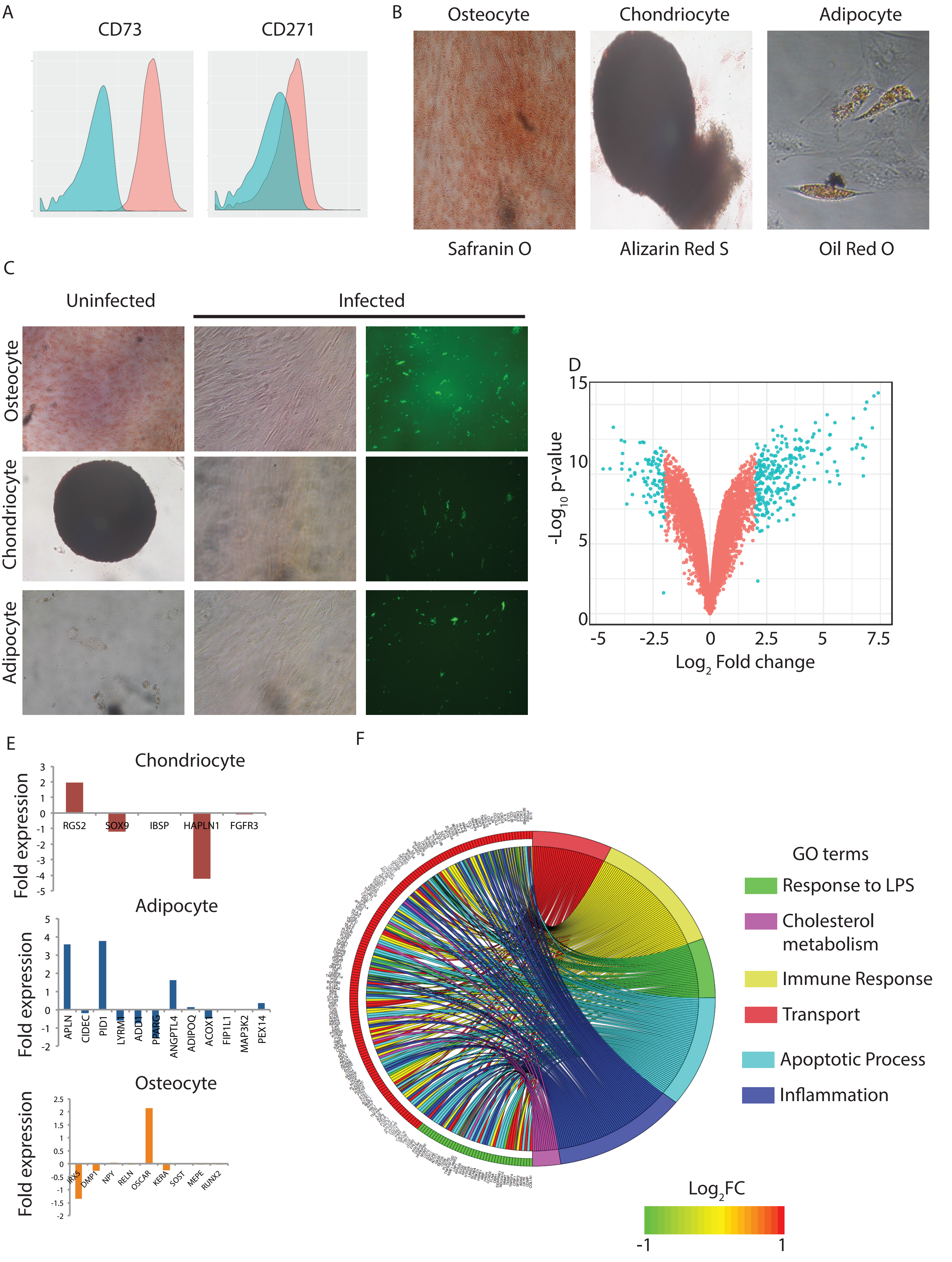

Supplementary figure S2

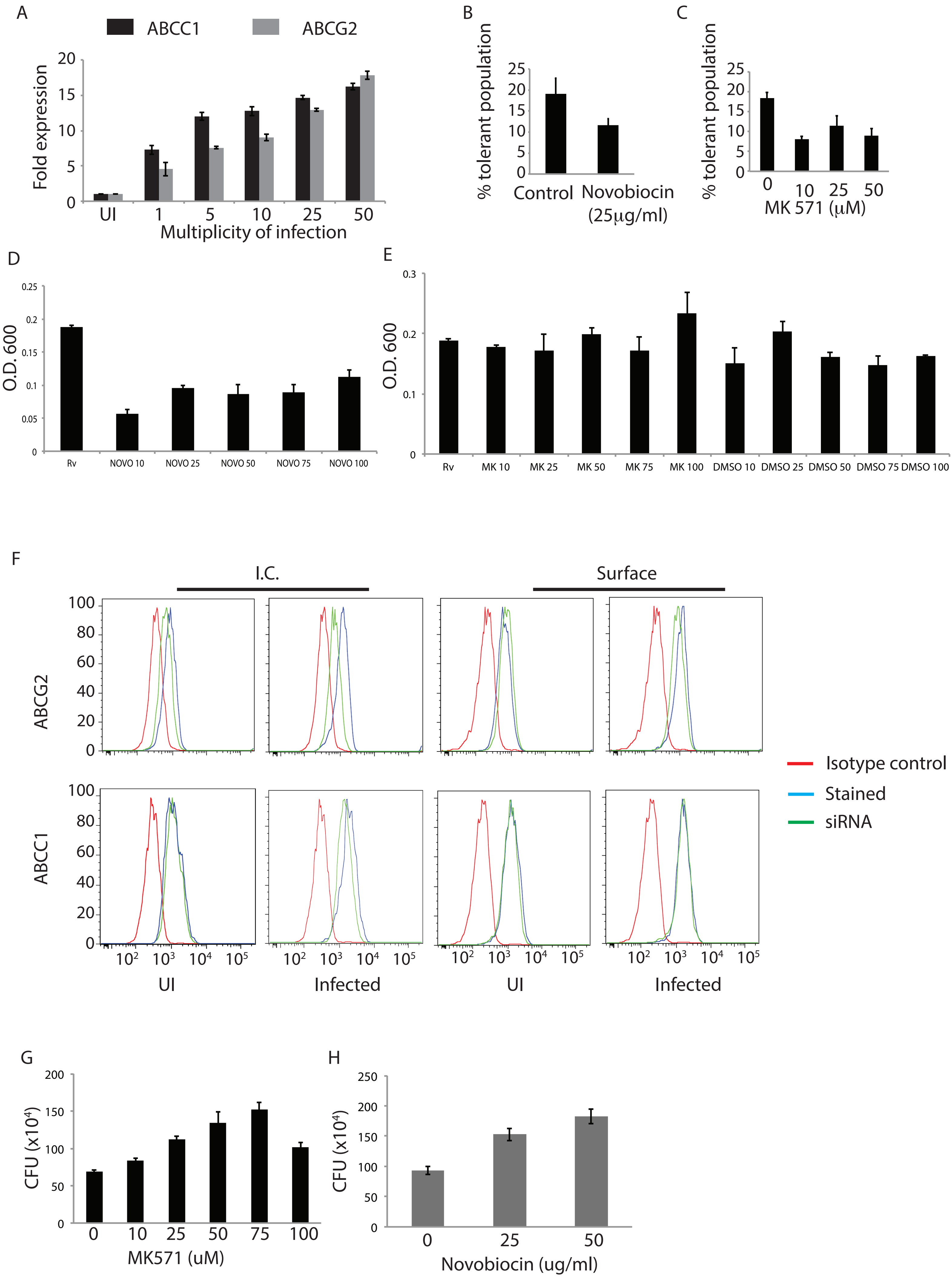

Supplementary figure S3

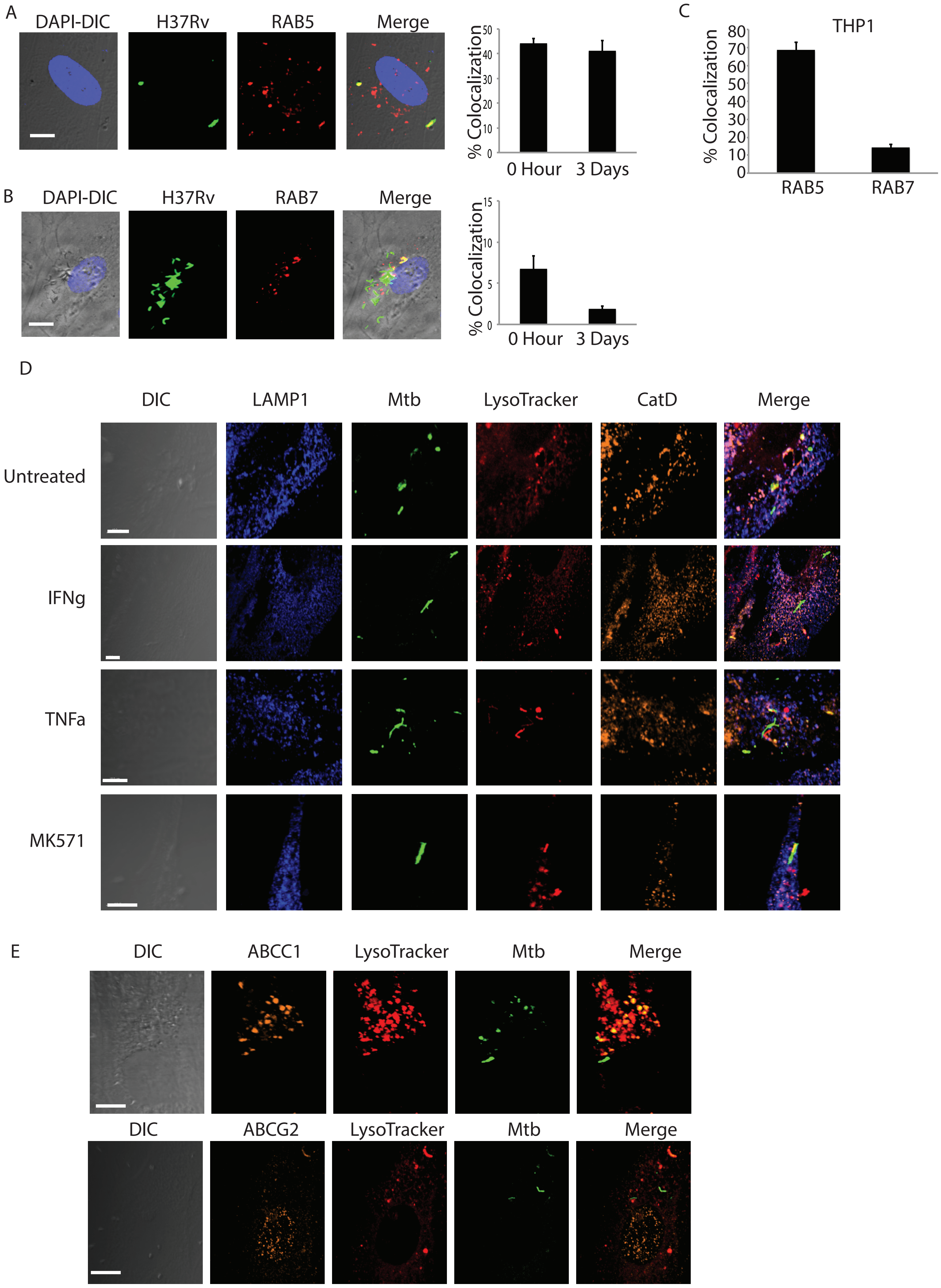

Supplementary figure S4

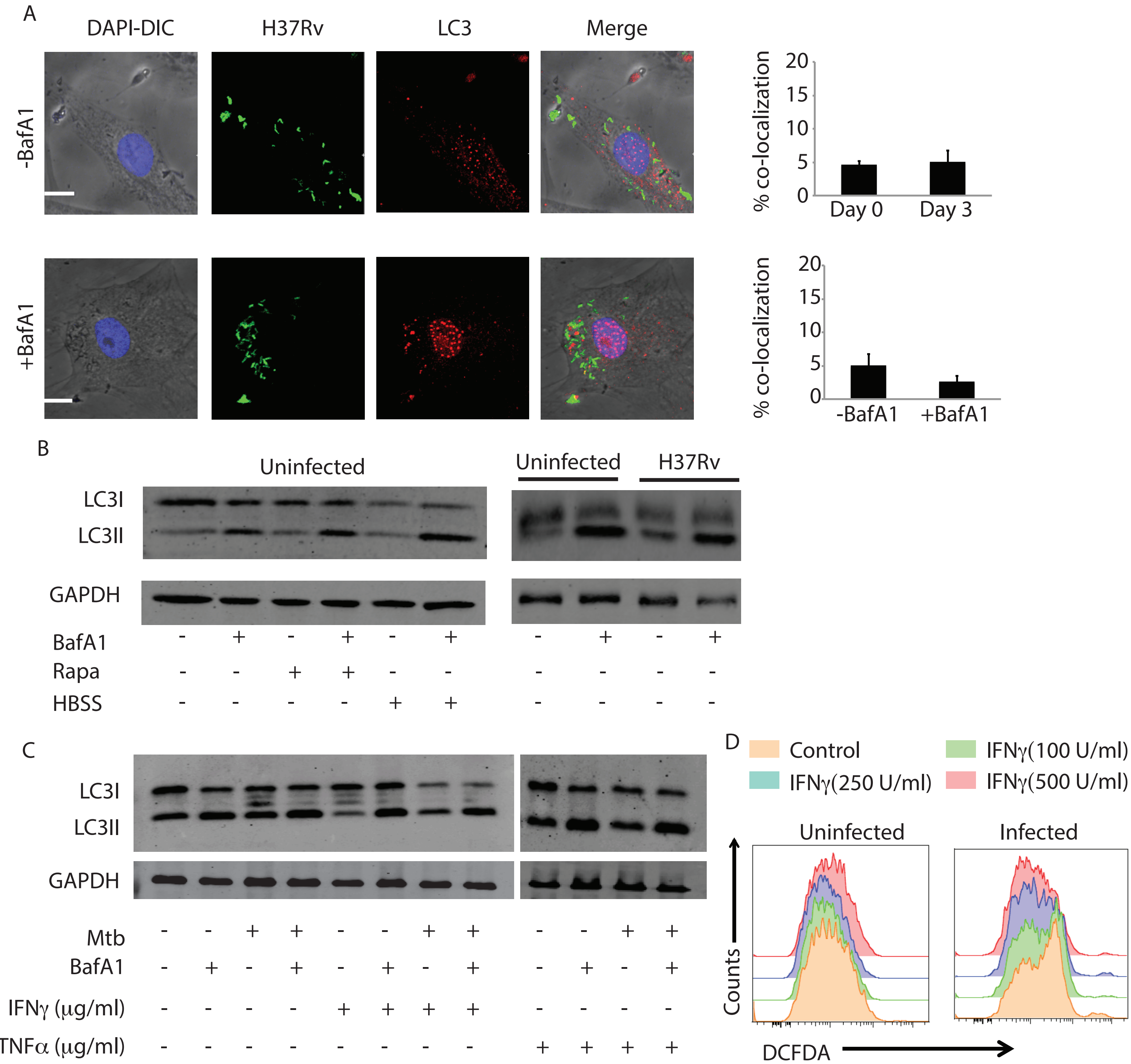

Supplementary figure S5

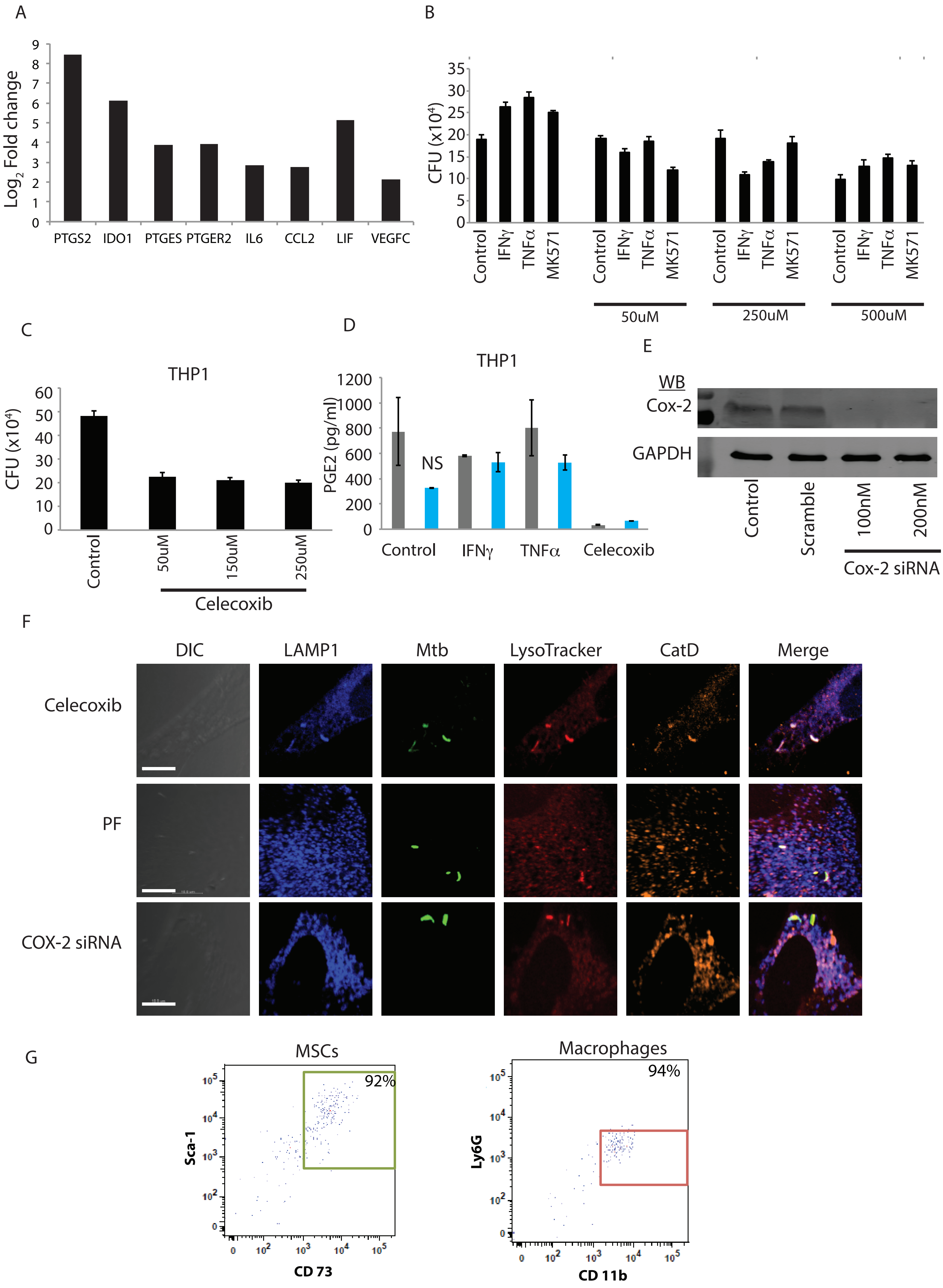
